# Inhibition of PLK1-dependent EBNA2 phosphorylation promotes lymphomagenesis in EBV-infected mice

**DOI:** 10.1101/2021.03.29.437455

**Authors:** Xiang Zhang, Patrick Schuhmachers, André Mourão, Piero Giansanti, Anita Murer, Sybille Thumann, Cornelia Kuklik-Roos, Sophie Beer, Stefanie M. Hauck, Wolfgang Hammerschmidt, Ralf Küppers, Bernhard Kuster, Monika Raab, Klaus Strebhardt, Michael Sattler, Christian Münz, Bettina Kempkes

## Abstract

While Epstein-Barr virus (EBV) establishes a life-long latent infection in apparently healthy human immunocompetent hosts, immunodeficient individuals are at particular risk to develop lymphoproliferative B cell malignancies caused by EBV. A key EBV protein is the transcription factor EBV nuclear antigen 2 (EBNA2), which initiates B cell proliferation. Here, we combine biochemical, cellular and in vivo experiments demonstrating that the mitotic polo-like kinase 1 (PLK1) binds to EBNA2, phosphorylates its transactivation domain and thereby inhibits its biological activity. EBNA2 mutants that impair PLK1 binding or prevent EBNA2 phosphorylation are gain-of-function mutants. They have enhanced transactivation capacities, accelerate the proliferation of infected B cells and promote the development of monoclonal B cell lymphomas in infected mice. Thus, PLK1 coordinates the activity of EBNA2 to attenuate the risk of tumor incidences in favor of the establishment of latency in the infected but healthy host.

## INTRODUCTION

Epstein-Barr virus (EBV) is associated with multiple malignancies including B cell lymphomas and B lymphoproliferative diseases (Farrell, 2019; Longnecker *et al*, 2013; Shannon-Lowe & Rickinson, 2019). More than 90% of the world population are infected with EBV. Based on an intimate virus-host interaction, the virus establishes a latent infection in resting memory B cells (Thorley-Lawson & Gross, 2004). Infected B cells are activated, enter the cell cycle and proliferate and maintain the viral circular genomes in a process that is strictly coordinated with the cell cycle of the host cell. While entering the memory B cell compartment, viral gene expression is gradually silenced to evade host immune surveillance. Reactivation of viral replication and virus production occurs occasionally (Munz, 2019).

EBV nuclear antigen 2 (EBNA2) is a key transcription factor that initiates and maintains the expression of viral and cellular target genes which are critical for the growth transformation of B cells by EBV (Kempkes & Ling, 2015; West, 2017). In the nucleus, EBNA2 associates with cellular proteins to execute its function. The B cell-specific transcription factor EBF1 (Friberg *et al*, 2015; Glaser *et al*, 2017; Lu *et al*, 2016) and CBF1/RBPJ, the major downstream effector of NOTCH signaling, serve as DNA anchors for EBNA2 (Farrell *et al*, 2004; Henkel *et al*, 1994; Hsieh & Hayward, 1995; Ling & Hayward, 1995). The C-terminal acidic transactivation domain (TAD) of EBNA2 recruits components of the pre-initiation complex and histone acetylases and also contributes to ATP-dependent chromatin remodeling by interacting with hSNF5/Ini (Cohen, 1992; Cohen & Kieff, 1991; Cohen *et al*, 1991; Kwiatkowski *et al*, 2004; Tong *et al*, 1995a; Tong *et al*, 1995b; Tong *et al*, 1995c; Wang *et al*, 2000). The TAD of EBNA2 is intrinsically unstructured. In complex with the histone acetylase CBP or the TFIIH subunit Tfb1, a 9-residue amphipathic α-helix within this TAD is formed (Chabot *et al*, 2014). Cellular proteins may control EBNA2 transcriptional activities by protein-protein interactions or modification. The MYND domain of the repressor BS69 can bind as a dimer to the EBNA2 TAD and its flanking regions and attenuates EBNA2 activity as well as transformation efficiency (Harter *et al*, 2016; Ponnusamy *et al*, 2019).

In search for additional cellular factors controlling EBNA2 function, we have performed a label-free mass spectrometry-based quantification of cellular proteins in EBNA2 immuno-precipitates and found polo-like kinase 1 (PLK1). PLK1 is a serine/threonine-protein kinase that controls G2/M transition and progress through mitosis and cytokinesis in a tightly controlled order to secure genomic stability of the dividing cell. Through phosphorylation of specific substrates, PLK1 promotes activation of the mitotic driver Cyclin B1/CDK1 in the late G2-phase, triggering prophase onset (Gheghiani *et al*, 2017; Nakajima *et al*, 2003; Watanabe *et al*, 2005). PLK1 recognizes its substrates by a conserved C-terminal polo-box domain (PBD). Frequently, PBD binds to pre-phosphorylated epitopes generated by the mitotic CDK1 kinase or other proline-directed kinases like MAPK (Elia *et al*, 2003a; Elia *et al*, 2003b; Lowery *et al*, 2005). This process is referred to as non-self-priming (Barr *et al*, 2004; Lee *et al*, 2014). Since primary tumors from various tissues express elevated levels of PLK1 and this high-level expression frequently correlates with a poor prognosis (Wolf *et al*, 1997), PLK1 is a potential oncotarget for molecular cancer therapy and a prognostic marker (Rodel *et al*, 2020; Rosenblum *et al*, 2020; Yuan *et al*, 1997). Preclinical and clinical studies currently test potential clinical indications for small molecule or siRNA-based PLK1 inhibitors (Liu *et al*, 2017). However, there is now increasing evidence that PLK1 can also act as a tumor suppressor when expressed in the context of specific tumor types. Thus there is a strong need to define the exact molecular features of tumors that should be treated with PLK1 inhibitors (de Carcer, 2019).

Here, we show that PLK1 directly binds to EBNA2. EBNA2/PLK1 complex formation is strongly enforced by EBNA2 residue S379 phosphorylation catalyzed by the mitotic Cyclin B/CDK1 complex. PLK1 phosphorylates the C-terminal transactivation domain of EBNA2 and attenuates its activity. EBNA2 mutants that lack either the main PLK1 docking site or the two phosphorylation sites are gain-of-function mutants that promote lymphoma incidence in EBV-infected humanized mice. This indicates that PLK1 acts as a tumor suppressor in EBV-driven carcinogenesis.

## RESULTS

### Cyclin B/CDK1 primes EBNA2 residue S379 for Polo-like kinase 1 (PLK1) binding

To identify potential EBNA2 interacting cellular proteins, we transfected EBV-negative DG75 B cells with HA-tagged EBNA2 expression constructs or the corresponding HA-expression vector and performed immunoprecipitations with HA-specific antibodies. Tryptic peptides of these immunoprecipitates were analyzed and quantified by label-free mass spectrometry. Polo-like kinase 1 (PLK1) was one of 19 proteins that were significantly enriched in EBNA2 co-immunoprecipitates (Table EV1). As expected, the EBNA2 DNA anchor protein CBF1/RBPJ was one of these proteins, demonstrating that the experimental approach was valid. Importantly, PLK1 and EBNA2 specifically co-immunoprecipitated from whole cell extracts of EBV-infected B cells in which both proteins are expressed at endogenous physiological levels (Fig. 1).

**Figure 1.**
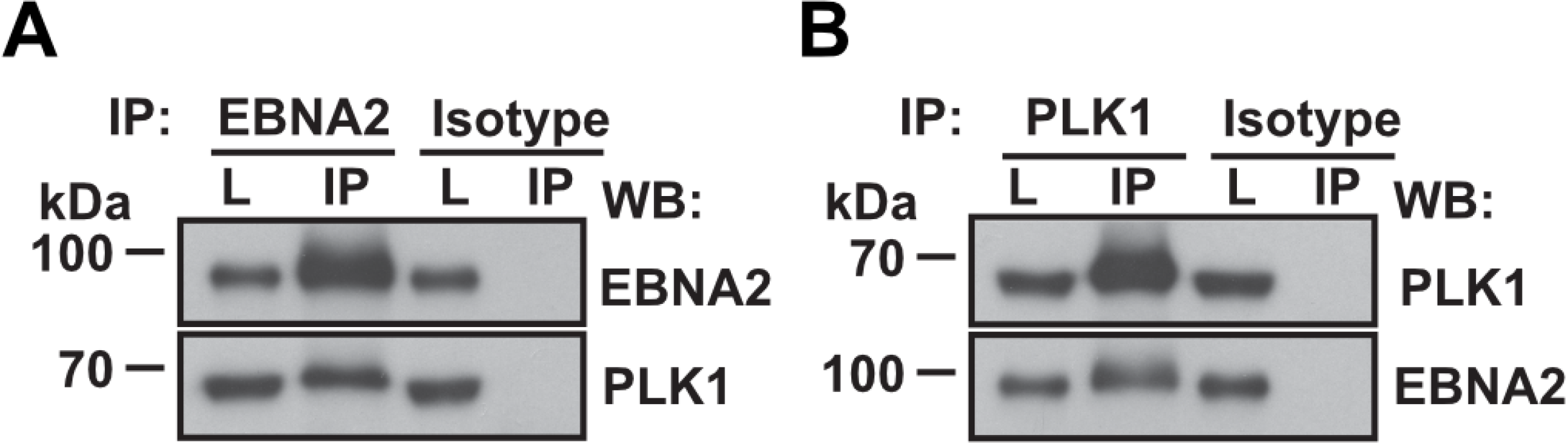
EBNA2 and PLK1 complexes in EBV infected B cell lines. Co-immunoprecipitation (IP) of EBNA2 (A) and PLK1 (B) using EBNA2-, PLK1-specific antibodies, or isotype-matched control antibodies performed on total cell lysates (L) of EBV immortalized B cells.

To identify PLK1 docking sites within EBNA2, EBNA2 deletion fragments were generated and tested for binding to PLK1 by transfection and co-immunoprecipitation (Fig. 1A, B, C). The smallest EBNA2 fragment that efficiently bound PLK1 mapped to region 342-474. Sub-fragments of either 327-487 or 342-474 showed some residual binding but none of these regions conferred binding efficiencies similar to the precursor.

PLK1 frequently binds to phosphorylated substrates that are primed by cellular kinases like Cyclin B/CDK1. These substrates share a consensus motif [Pro/Phe]-[Φ/Pro]-[Φ]-[Thr/Gln/His/Met]-Ser-[pThr/pSer]-[Pro/X] (Φ represents hydrophobic and X represents any residue) specific for binding to the Polo-box binding domain (PBD) of PLK1 (Elia *et al.*, 2003a). Crystal structures of the PLK1 PBD in complex with peptides demonstrate how the positively charged groove of PBD docks onto negatively charged phosphopeptides of diverse substrates. EBNA2 exhibits three potential Cyclin B/CDK1 phosphorylation/ PBD docking sites located at residue T267, S379 and S470 (Fig. D). Mutants involving the respective residues were tested for PLK1 binding by transfection and co-precipitation. The mutation TSS377VAA impaired PLK1 binding dramatically while all other EBNA2 mutations did not affect PLK1 binding (Fig. 1E). To specify the contribution of EBNA2 residue S379 phosphorylation, a heptapeptide (PNTSSPS) and phospho-heptapeptide (PNTSpSPS) were tested for PBD (PLK1 residue 345-603) binding by isothermal titration calorimetry (ITC). The phosphorylated peptide bound PBD in a molar ratio of 1:1 whereas no interaction was detected using the unmodified peptide. The PNTSpSPS/ PBD interaction is micromolar with a dissociation constant of *K*_D_ = 8.19 μM (Fig. 1F). GST EBNA2 342-422 expressed and purified from bacterial extracts did hardly pull-down any PLK1 from cellular extracts. Phosphorylation of the EBNA2 protein produced in bacteria by Cyclin B/ CDK1, an enzyme not present in bacteria, strongly enhanced PLK1 binding. EBNA2 (342-422) S379A mutant did not bind PLK1 and phosphorylation by Cyclin B/CDK1 could not reconstitute binding (Fig. 1G). Also, as shown by transfection, EBNA2 S379A binding to PLK1 was severely impaired (Fig. 1H). We conclude that EBNA2 residue S379 is a PLK1 docking site primed by Cyclin B/CDK1 phosphorylation.

### The C-terminus of EBNA2 is the substrate of PLK1

In order to test if EBNA2 is phosphorylated by PLK1, EBNA2/PLK1 complexes were coprecipitated from cellular extracts of DG75 cells induced with doxycycline to express HA-EBNA2 (DG75^Dox HA-EBNA2^) (Fig. 3A). The precipitates were submitted to kinase assays either in the presence or absence of exogenous recombinant PLK1. Phosphorylation of EBNA2 was readily detected in the co-precipitates and was enhanced by the addition of exogenous recombinant PLK1 to the co-precipitates (Fig. 3B). EBNA2 phosphorylation was prevented upon addition of Volasertib, a PLK1 specific inhibitor, to the co-precipitates of EBNA2 and endogenous PLK1 (Fig. 3C), corroborating the evidence that the endogenous PLK1 trapped in the co-precipitate is the active kinase.

**Figure 3.**
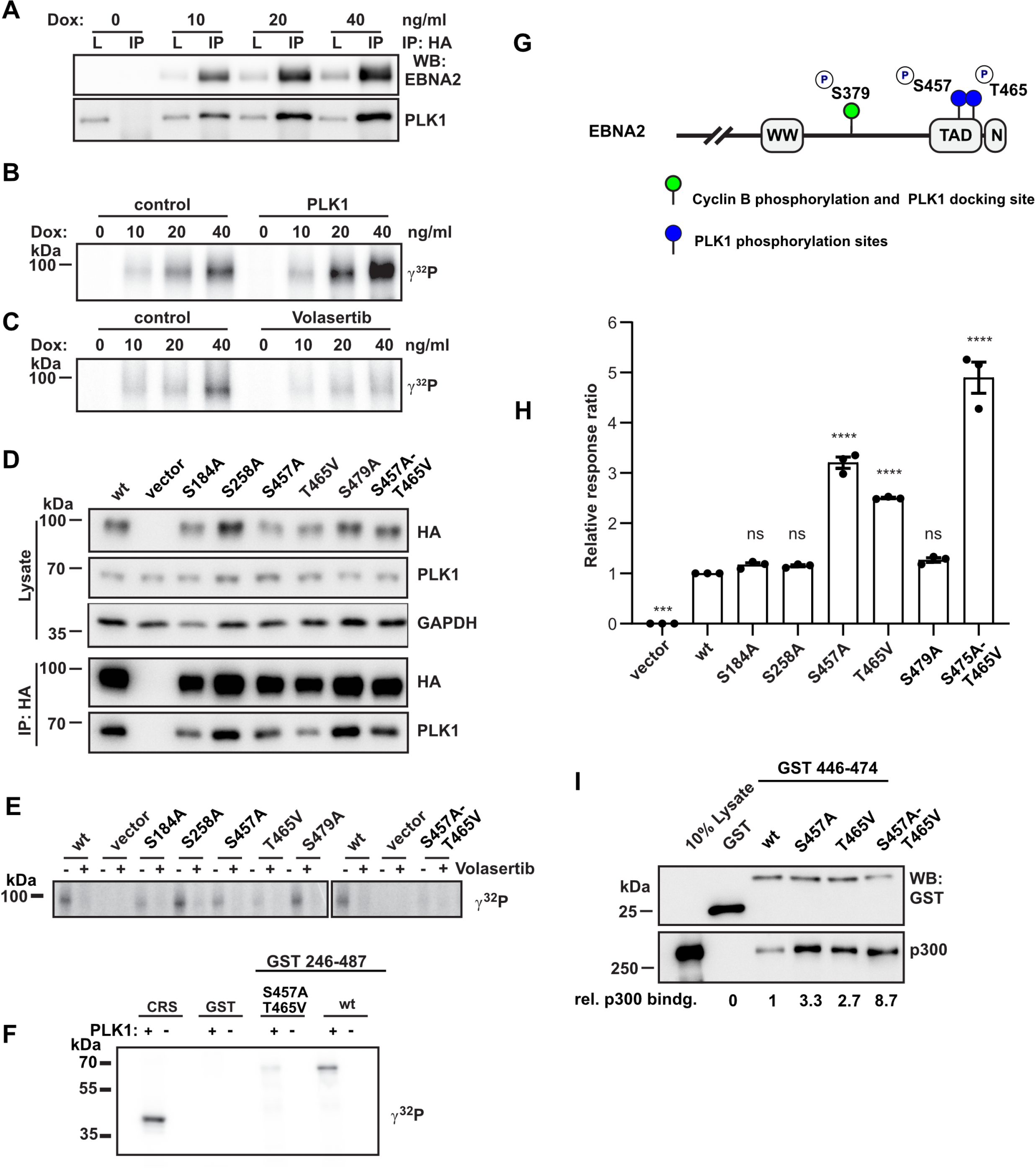
PLK1 phosphorylates the EBNA2 residue S457 and T465 within the C-terminal transactivation domain. (A) Doxycycline (Dox) induction of EBNA2 in DG75^Dox HA-EBNA2^ cells treated for 24 hours. Co-immunoprecipitates of HA-EBNA2 and endogenous PLK1 are visualized by Western blotting. (B) EBNA2/PLK1 co-precipitates were submitted to kinase reactions using [γ-^32^P] ATP in the absence (control) or presence of 50 ng recombinant active PLK1. (C) EBNA2 co-precipitates were submitted to kinase reactions in the absence (control) or the presence of the PLK1 inhibitor Volasertib (40 nM). (D) EBNA2 candidate phosphorylation mutants were expressed in DG75 B cells and tested for PLK1 binding by co-immunoprecipitations followed by Western blotting. (E) Immunoprecipitates were submitted to kinase reactions as in B, but samples were treated with Volasertib (40 nM) (+) or treated with solvent only (-). (F) GST EBNA2 fragment 246-487 wt and mutant S457A/T465V were treated with recombinant active PLK1 (+) in the presence of [γ-^32^P] ATP in vitro or left untreated (-). CRS, an artificial PLK1 test substrate (Yuan *et al*, 2002), was used as a positive control. (G) Schematic presentation of EBNA2 phosphorylation sites by CDK1 and PLK1. (H) HA-tagged EBNA2 candidate phosphorylation mutants were expressed in DG75 B cells and tested for activation of an EBNA2/CBF1 responsive promoter reporter luciferase plasmid. Activation of the reporter gene is shown as relative response ratio normalized to Renilla luciferase activity and shown relative to wt activity. (I) GST-pull down assay using GST-EBNA2 446-474 as a bait to purify cellular proteins from DG75 cells followed by Western blotting and quantification of signals obtained by GST and p300 specific antibodies. Relative binding affinities (rel. p300 bindg.) are normalized to the wt signal.

**Figure 4.**
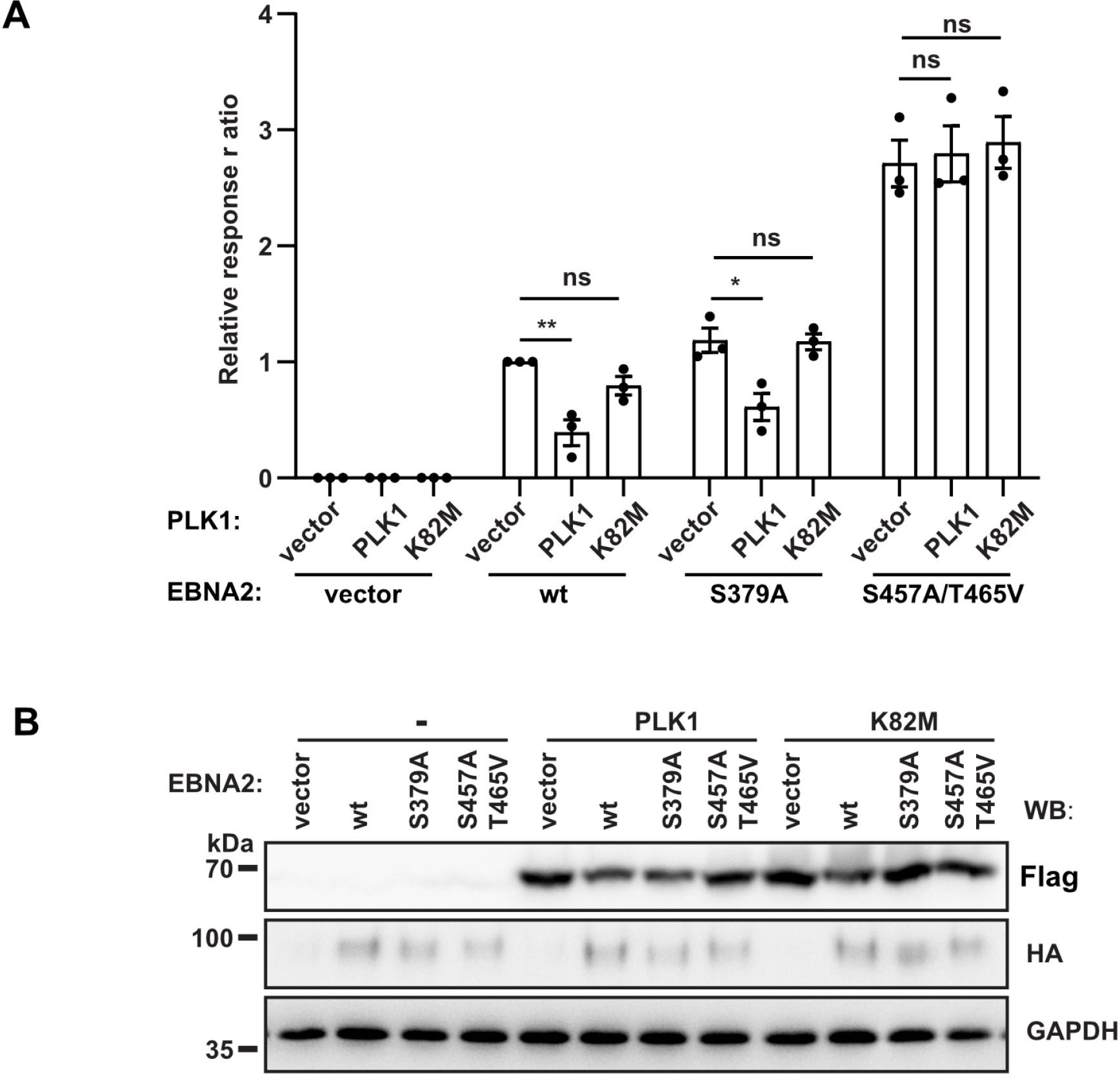
PLK1 inhibits EBNA2 activity by phosphorylating residues S457/T465. (A) Flag-tagged active (PLK1) and kinase inactive PLK1 (K82M) were co-expressed with HA-EBNA2 wt, docking site mutant (S379A) and phosphorylation mutant (S457A/T465V) and tested for activation of an EBNA2/CBF1 responsive promoter reporter. Activation of the reporter firefly luciferase gene is shown as relative response ratio normalized to Renilla luciferase activity and shown relative to EBNA2 wt activity. (B) 30 μg of cellular extracts that were produced for luciferase assays were tested for protein expression using Flag-, HA and GAPDH specific antibodies.

To identify the amino acid residues that are phosphorylated by PLK1, bacterially expressed EBNA2 was phosphorylated by active recombinant PLK1 or left untreated. The EBNA2 protein was digested by trypsin and endoproteinase Glu-C (V8 Protease) in parallel. Tryptic or V8 Protease derived peptides and phospho-peptides were identified by mass spectrometry (MS). Since neither tryptic nor V8 derived peptides covered the C-terminus of EBNA2 sufficiently, a subfragment (453-474) of EBNA2 flanked by arginine residues and expressed as a GST fusion protein was used for further tryptic digest and phosphopeptide mapping (Fig. EV1). Initially, 11 potential phosphorylation sites were found. Of these, 5 phosphorylation sites (S184, 258, 457, 479 and T465) were confidently localized (Fig. EV2) and 6 additional sites (T175, 178, 263, 267, 464 and S266) were ambiguously mapped. To test, if the 5 confident phosphorylation sites identified in vitro are relevant also in cells, EBNA2 mutants with a singular or combined mutations were generated and expressed in DG75 B cells. Immunoprecipitations and subsequent kinase assays based on endogenous PLK1 trapped in the precipitates were performed (Fig. 3D). None of the mutations severely impaired PLK1 binding but Volasertib prevented phosphorylation of all EBNA2 proteins. In the absence of Volasertib, phosphorylation of S184A, S258A and S479A was not reduced. Phosphorylation of the EBNA2 mutants S457A and T465V was impaired, and the combination of both mutations (S457A/T465V) abolished phosphorylation (Fig. 3E). To corroborate this finding GST EBNA2 246-487 either in wild-type or as S457A/T465V mutant was submitted to kinase assays using recombinant PLK1 in vitro. This experiment resulted in efficient phosphorylation of wild-type but not mutant EBNA2 S457A/T465V (Fig. 3F). We conclude that the PLK1 docking and the PLK1 phosphorylation site are two independent EBNA2 modules that can be dissected and analyzed by specific mutations (Fig. 3G).

To test the phosphorylation mutants for their biological activity, we used a luciferase reporter construct driven by an EBNA2 responsive artificial promoter that harbors a multimerized CBF1 binding site to recruit EBNA2 (Minoguchi *et al*, 1997). All mutants that carried either the S457A, the T465V mutation or both mutations showed enhanced transactivation potential, suggesting that PLK1 might negatively regulate EBNA2 activity (Fig. 2H). Since the C-TAD of EBNA2 is known to bind the histone acetylase and co-activator p300 (Chabot *et al.*, 2014), p300 binding to EBNA2 phosphorylation mutants was tested by GST-pulldown experiments. While p300 binding of the single EBNA2 phosphorylation mutants was strongly enhanced by approximately 2-3-fold, binding by the double mutant S457A/T465V was increased even 8-9-fold. Thus, enhanced p300 binding of EBNA2 phosphorylation mutants correlates well with improved transactivation activity. This finding suggests that PLK1 phosphorylation hinders p300 recruitment to EBNA2, thereby inhibiting EBNA2 transactivation (Fig. 3I).

**Figure 2.**
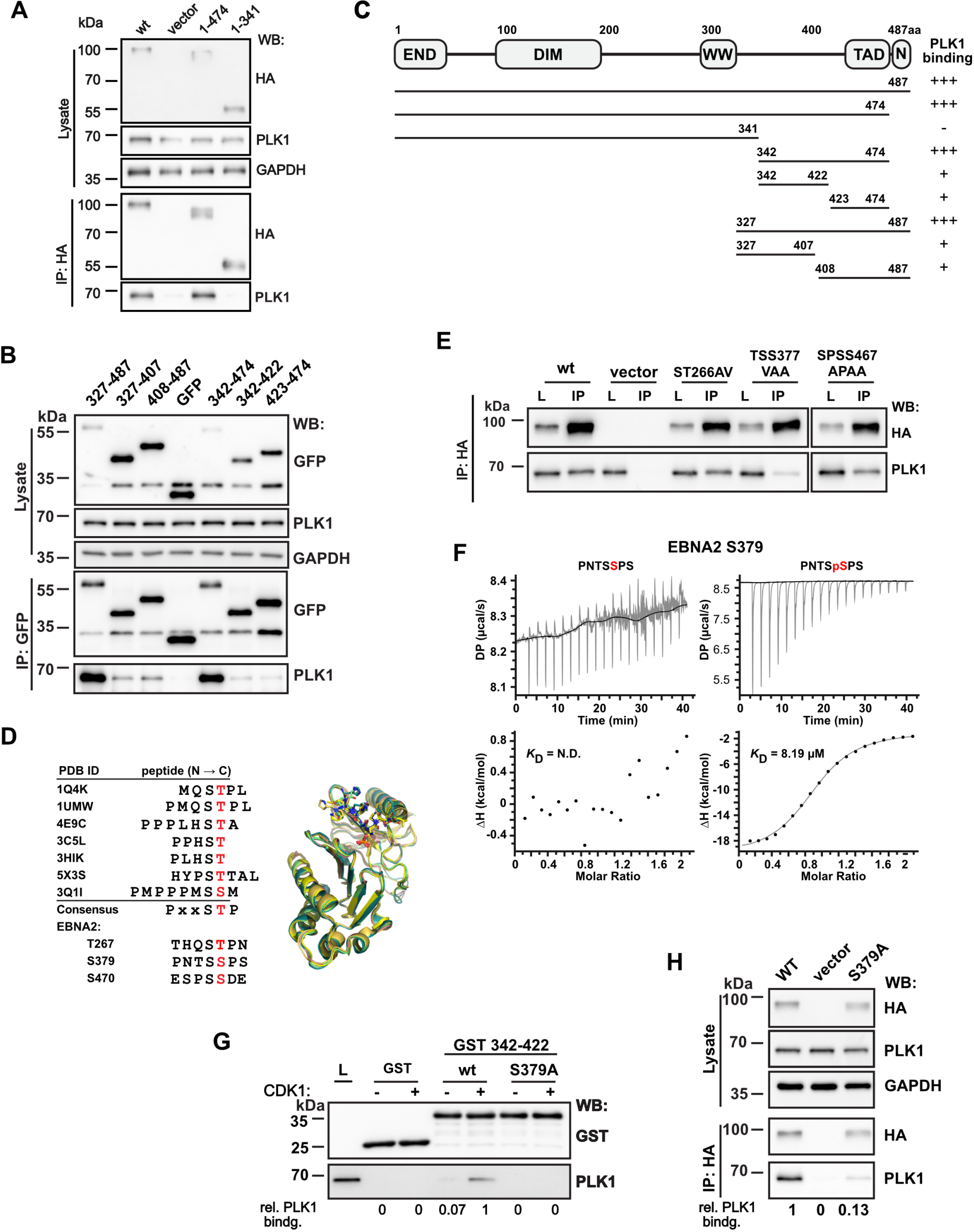
EBNA2/PLK1 phosphorylation-dependent complex formation. Transfection and immunoprecipitation of (A) HA-tagged or for smaller fragments (B) GFP-tagged EBNA2 fragments to co-precipitate endogenous PLK1. Total protein lysates and immunoprecipitates (IP) were analyzed by Western blotting (WB). (C) Schematic outline of EBNA2, its dimerization domains (END, DIM), the region used by CBF1 to recruit EBNA2 to DNA (WW), the C-terminal transactivation domain (TAD), the nuclear localization signal (N) and the EBNA2 fragments used to map the PLK1 docking site (UniProt ID: P12978.1). The panel on the right summarizes the results of the co-immunoprecipitations in (A) and (B). (D) Multiple sequence alignment (left) and superposition (right) of several phosphopeptides present in published crystal structures of PLK1 PBD (phosphorylated residues stained red). Crystal structures of the PBD in complex with peptides show that the positively charged groove of PBD docks in a similar mode to the negatively charged phosphopeptides. References for PDB ID: 1Q4K (Cheng *et al*, 2003), 1UMW (Elia *et al.*, 2003a), 4E9C (Śledź *et al*, 2012), 3C5L (Yun *et al*, 2009), 3HIK (Yun *et al.*, 2009), 5X3S (Lee *et al*, 2018), 3Q1I (Pavlovsky *et al*, 2012). Potential residues of EBNA2, which might be a PBD docking site, are listed below. (E) Immunoprecipitation of HA-tagged EBNA2 mutant ST266AV, TSS377VAA and SPSS467APAA using HA-specific antibodies. (F) ITC thermogram of PLK1 PBD titrated with the peptide PNTSSPS or the phosphopeptide PNTSpSPS of EBNA2. (G) GST-pulldown of PLK1 from total cellular extracts using Cyclin B/CDK1 phosphorylated GST-EBNA2 region 342-422. (H) Co-immunoprecipitation of transfected EBNA2 wt or S379A and endogenous PLK1.

To directly test if PLK1 inhibits EBNA2 functions, PLK1 and EBNA2 were co-expressed and luciferase reporter assays were performed. EBNA2 wt activity was significantly reduced by co-expression of PLK1 but not affected by the kinase dead K82M PLK1 mutant. The activity of the docking site mutant S379A was weakly impaired by co-expression of PLK1 suggesting that residual binding activity, as demonstrate in Fig. 2H, might still recruit PLK1 activity. Importantly, the transactivation capacity of the phosphorylation mutant S457A/T465V was not affected by PLK1 co-expression. Since the kinase dead PLK1 mutant did not impair the transactivation capacity of any EBNA2 protein, we conclude that PLK1 binding is necessary but not sufficient to inhibit EBNA2 and the phosphorylation of EBNA2 by PLK1 is required to inhibit EBNA2.

In summary, PLK1 phosphorylates S457 and T465 within the TAD of EBNA2 to attenuate its activity and to impair p300 binding. PLK1 uses S379A as a phosphorylation-dependent docking site that can be primed by Cyclin B/CDK1.

### EBV strains expressing EBNA2 mutants deficient for PLK1 docking or phosphorylation are gain-of-function mutants in B cell immortalization assays

To test EBNA2 mutants defective for PLK1 binding or phosphorylation by PLK1 for their B cell immortalization potential, three new viral mutants based on the viral backbone of the recombinant EBV strain p6008 (Mrozek-Gorska *et al*, 2019; Pich *et al*, 2019) were generated. Carboxyl-terminally HA-tagged EBNA2 (EBV wt), HA-tagged EBNA2 S379A (EBV S379A), and HA-tagged EBNA2 S457A/T468V (EBVS457A/T465V) were inserted into the backbone of the recombinant EBV strain p6008 by homologous recombination using a two-step positive-negative selection protocol, also called recombineering (Wang *et al*, 2009). The integrity of the viral genomes was controlled by restriction digest and sequencing (Fig. EV3). Infectious viruses were produced in HEK293 cells. Primary B cells from three unrelated donors were infected. During the first 6 days post-infection, the proliferation rates of EBV wt, EBV S457A/T465V or EBV S379A infected B cells were determined. The phosphorylation mutant S457A/T465V proliferated the fastest, followed by S379A and wt infected cells (Fig.5A and Fig. EV3C, D). Long-term cultures could be established and the proliferation of these lymphoblastoid cell lines (LCLs) was characterized by counting the cells daily (Fig. 5B). The proliferation rates of both long-term cultures infected with mutant viruses were higher than EBV wt infected cells. EBNA2 expression levels in these long-term cultures were similar for wt and S379A but less pronounced for the S457A/T465V mutant. LMP1 expression varied between the cell lines obtained from different donors but was consistently elevated in EBV S457A/T465V infected cell lines compared to S379A and EBV wt infected B cells. MYC expression was seen in all infected cultures and MYC levels were not affected by individual virus variants (Fig. 5C). In summary, both mutants were gain-of-function mutants with respect to proliferative capacities of the infected cell cultures and LMP1 activation. In comparison, these features were more pronounced in EBV S457A/T465V than in S379A infected B cells.

**Figure 5.**
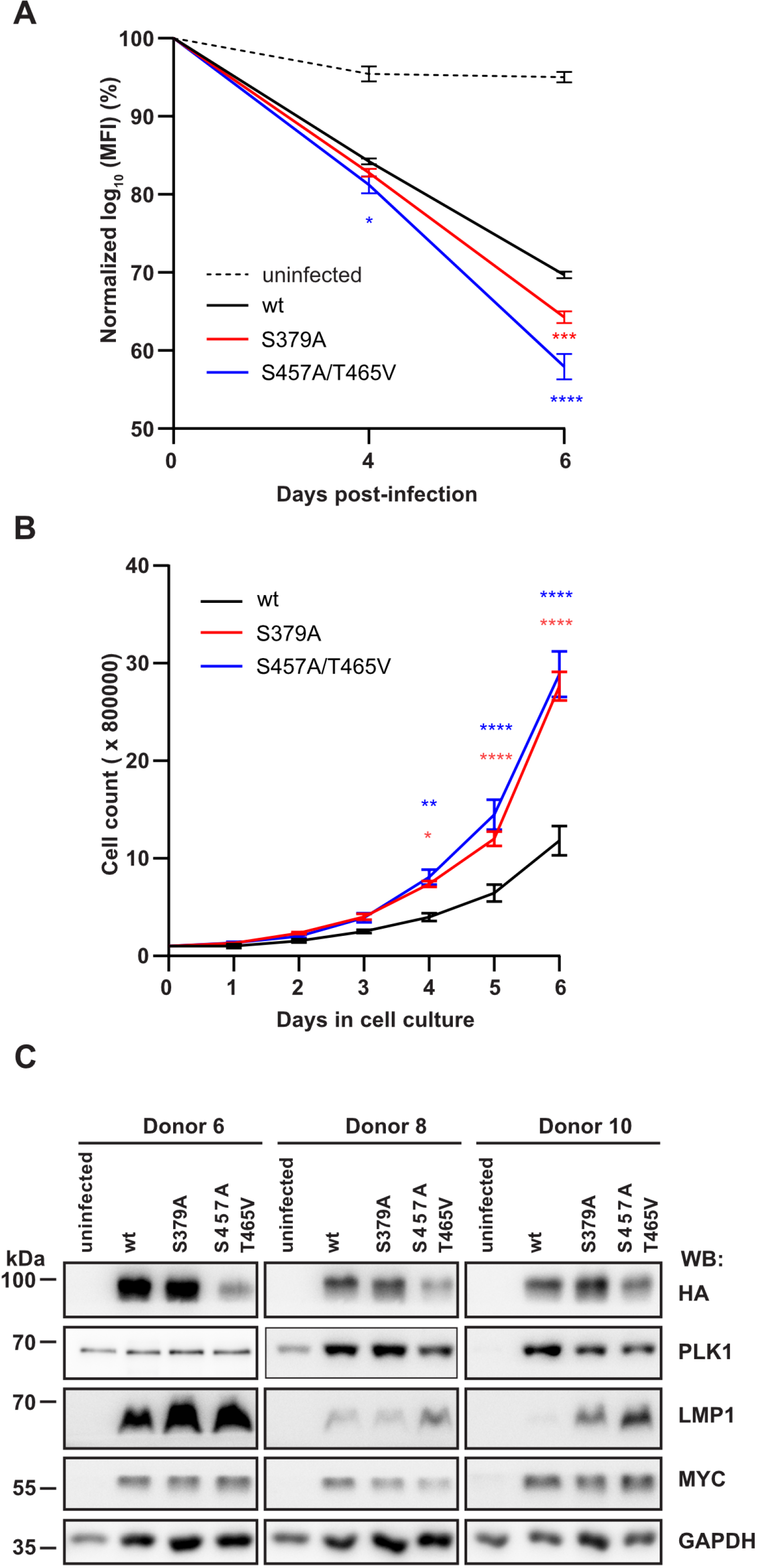
EBV strains expressing EBNA2 mutants deficient for PLK1 docking or phosphorylation accelerate B cell proliferation and express elevated levels of the viral LMP1 protein. (A) Primary B cells were stained with cell trace violet before they were infected and cell cultures were started. Loss of the fluorescent cell tracking dye was recorded by flow cytometry to monitor proliferation on day 0, 4 and 6 post-infection. Mean values for 3 biological replicates, standard deviation and significance of changes in signal loss of mutants compared to wild-type controls (p-value, *: p < 0.05; ****: p < 0.0001). (B) Long-term growth transformed B cell cultures infected with recombinant EBV mutants were seeded at a starting concentration of 2 x 10^5^ cells per ml. Data were presented as the mean ± S.E.M. of n = 4 biological replicates. Statistical significance was tested by two-way ANOVA followed by a Tukey’s multiple comparison test (*: P < 0.05, **: P < 0.01, ****: P < 0.0001, vs WT). (C) Expression of EBNA2, LMP1, PLK1, MYC, and GAPDH was analyzed by Western blotting and immunostaining of whole cell extracts of cell lines established from 3 individual donors.

### Enhanced frequencies of B cell lymphomas in humanized mice infected with EBV S379A or EBV S457A/T465V

Since EBV does not infect murine B cells, tumor development and immune control were studied in humanized mice. Immunodeficient NSG (NOD scid γ_c_^−/−^) mice can be engrafted with CD34^+^ human hematopoietic progenitor cells to develop human immune system components that can be readily infected with human lymphotropic pathogens, including EBV. These chimeric humanized mice were used previously to study EBV-induced tumor formation after EBV infection and anti-viral immune mechanisms against EBV in vivo (Strowig *et al*, 2009b). Hence, we infected three cohorts (designated experiment 1, 2 and 3) of humanized mice intraperitoneally with 10^5^ Green Raji Units (GRU) of EBV and analyzed viral blood loads, CD8^+^ and CD4^+^ T cell composition as well as activation in their blood in weekly intervals. Five weeks post-infection, all EBV-positive mice were sacrificed to analyze tumor development (Fig. 6A). In two out of three independent experiments, infection with EBV S457A/T465V caused increased mortality rates compared to EBV wt or S379A infected animals (66% survival in the second and 20% in the third experiment; Fig. 6B). In addition, mice that were infected with either of the two EBV mutants presented with higher incidence of tumor development in spleen, peritoneal cavity or lymph nodes (23% for WT, 36% for S379A, and 44% for S457A/T465V; Fig. 6C). Viral loads in the spleen of mice infected with mutants were slightly elevated 5 weeks post-infection when mice were sacrificed (Fig. 6D). In blood, viral loads of EBV wt and EBV S379A infected mice gradually increased until week 3 and remained at similar levels until the end of the experiment. In contrast, viral loads in mice within the EBV S457A/T465V mutant group reached higher levels already two weeks after infection and started to decrease at 4 weeks post-infection (Fig. 6E).

**Figure 6.**
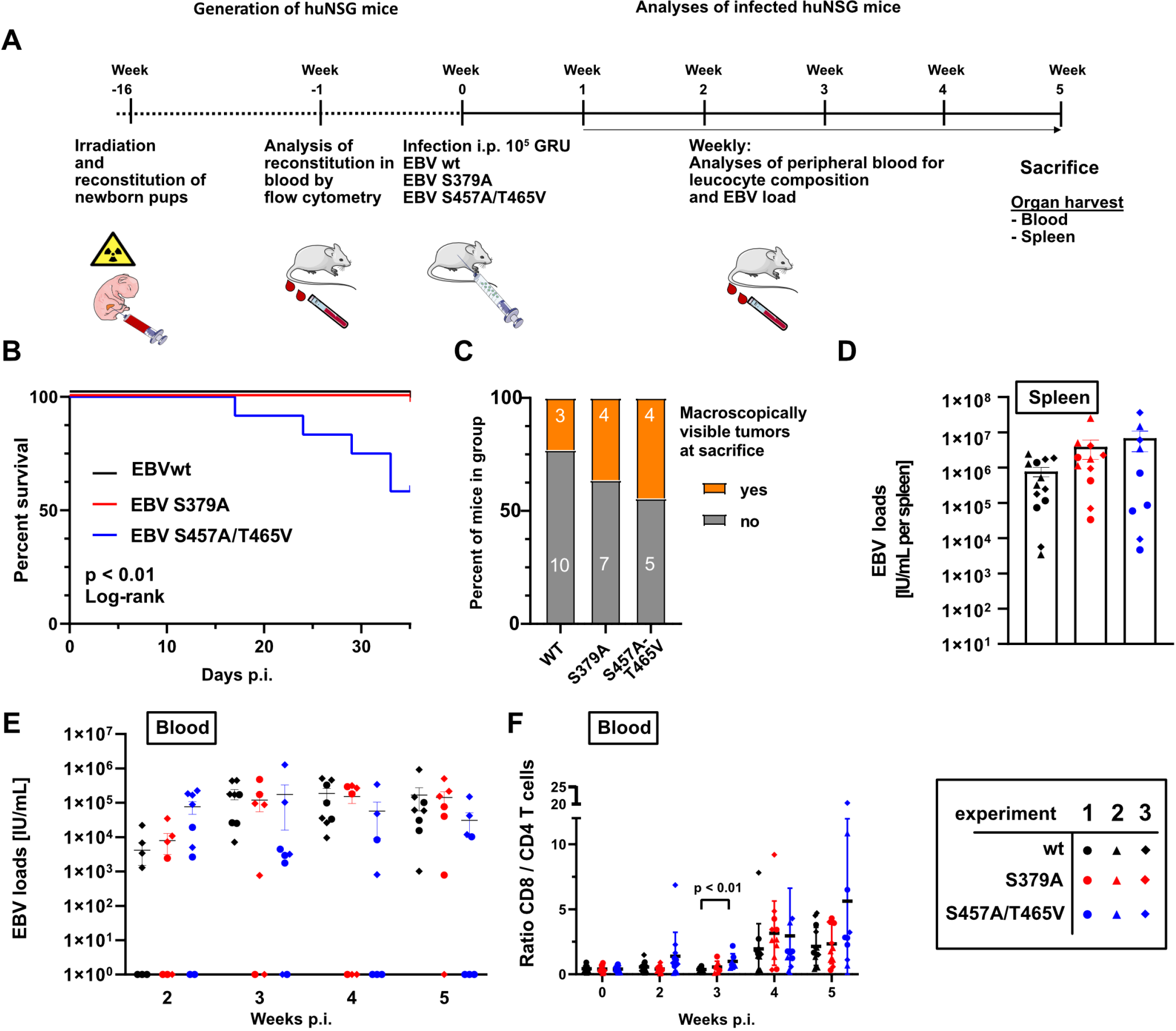
EBV strains carrying EBNA2 mutants with diminished PLK1 binding or resistant to PLK1 phosphorylation cause lymphomas in humanized mice more frequently than EBV wt. (A) Experimental set-up. (Images of animals, syringes and blood collection tubes are derived from Servier Medical Art.) (B) Survival curve of humanized mice infected with 10^5^ GRU EBV wt (n = 13), EBV S379A (n = 11) or EBV S457A/T465V EBV (n = 9). Log-rank test. (C) Percentage of mice having macroscopically visible tumors at the day of sacrifice. Numbers within bars indicate the total number of mice with or without tumors in the respective groups. Fisher’s Exact test. (D) Viral loads in spleen of infected humanized mice at the day of sacrifice. Error bars indicate mean ± SEM. EBNA2 WT EBV: n = 13; EBNA2 S379A EBV: n = 11; EBNA2 S457A T465V EBV: n = 9. (E) Development of viral loads in blood of infected mice over a period of five weeks. Error bars indicate mean ± SEM. Mann-Whitney U test. (F) Flow-cytometric analyses of CD8^+^/CD4^+^ T cell ratios in the blood of infected mice over a period of five weeks. Error bars indicate mean ± SD.

Extensive CD8^+^ T cell expansion and activation in blood is a common trait marking EBV infection in humanized mice. It usually follows, with a delay of about a week, rising viral loads (Chatterjee et al., 2019; Strowig et al., 2009) and correlates with these (Caduff *et al*, 2020; Zdimerova *et al*, 2020). As CD8^+^ T cells expand more strongly than CD4^+^ T cells upon EBV infection, a rising CD8^+^ to CD4^+^ T cell ratio indicates the extensive proliferation of CD8^+^ T cells.

Three weeks post-infection the mean CD8^+^/CD4^+^ T cell ratios in mice infected with EBV S457A/T465V (0.99) were significantly higher than in EBV wt (0.37) infected mice (Fig. 6F), while CD8^+^/CD4^+^ T cell ratios of EBV wt infected animals increased later, at 4 weeks postinfection (Fig. 6F).

CD8 and CD4 T cell populations infected with mutant virus started to expand already 2 weeks post-infection (Fig. EV4 A, B). Also, the relative fraction of CD8^+^ T cells expressing the activation marker HLA-DR increased from week 2 post-infection, thus earlier than the control EBV wt infected group (Fig. EV4 C, D). A similar trend could be observed for mice infected with the EBNA2 S379A mutant. Interestingly, the EBV S457A/T465V mutant more strongly induced the activation of CD4^+^ T cells in the blood of mice within 5 weeks of infection compared to the EBV wt infected group (Fig. EV4 D). In contrast to blood, the fractions of CD8^+^ and CD4^+^ T cells in the spleen of infected animals did not differ between the groups (Fig. EV4 E) but a higher fraction of CD8^+^ and a significantly increased fraction of CD4^+^ T cells were HLA-DR-positive and thus activated in EBV S457A/T465V infected animals (Fig. EV4 F, G). The development of memory T cells, i.e. effector memory (Tem), central memory (Tcm) or terminally differentiated subsets that re-express CD45RA (Temra), however, was not influenced by EBNA2 mutations (Fig. EV4 H, I). In summary, EBV S379A and EBV S457A/T465V infections developed more rapidly in humanized mice, resulting in earlier immune responses, increased lymphomagenesis and more frequent demise of the mice.

### Tumors of EBV S379A or EBV S457A/T465V infected mice frequently present as monoclonal B cell lymphomas

The diversity of immunoglobulin gene rearrangements and sequences can be used to define the clonal composition of a given B cell population (Kuppers *et al*, 1993; Rajewsky, 1996). In experiment 2, 1 out of 5 EBV wt, 2 out of 4 EBV S379A and 2 out of 4 EBV S457A/T465V infected mice developed tumors. The clonality of macroscopically visible tumors detected in experiment 2 was assessed by PCR of rearranged IgH V genes, using subgroup specific IgH V primers together with an IgH J primer mix in separate reactions, followed by Sanger sequencing of amplificates obtained (Table 1). Polyclonal B cell expansions were found in the splenic tumors of EBV wt (#13) and in one splenic tumor of EBV S379A (#23). In all of the four analyzed tumor bearing mice infected with mutant EBV, monoclonal B cell populations were identified, sometimes on a background of remaining polyclonal B cells. In mouse #23, the spleen harbored a polyclonal B cell population, while the peritoneum showed a monoclonal B cell expansion. This analysis validates that the tumors developing in EBNA2 mutant mice are indeed monoclonal, and hence bona fide malignant B cell lymphomas. In all five monoclonal lymphomas, productive in-frame IgH V genes were obtained. Three of the samples showed additional out-of-frame rearrangements, which likely represent the second IgH alleles of the monoclonal lymphomas defined by the in-frame rearrangements. Since humanized NOD scid γ_c_^−/−^ mice in the experimental setup of our study cannot generate germinal centers we did not expect somatic hypermutation to occur. Indeed, somatic mutations were not observed in any specimen.

**Table 1.**
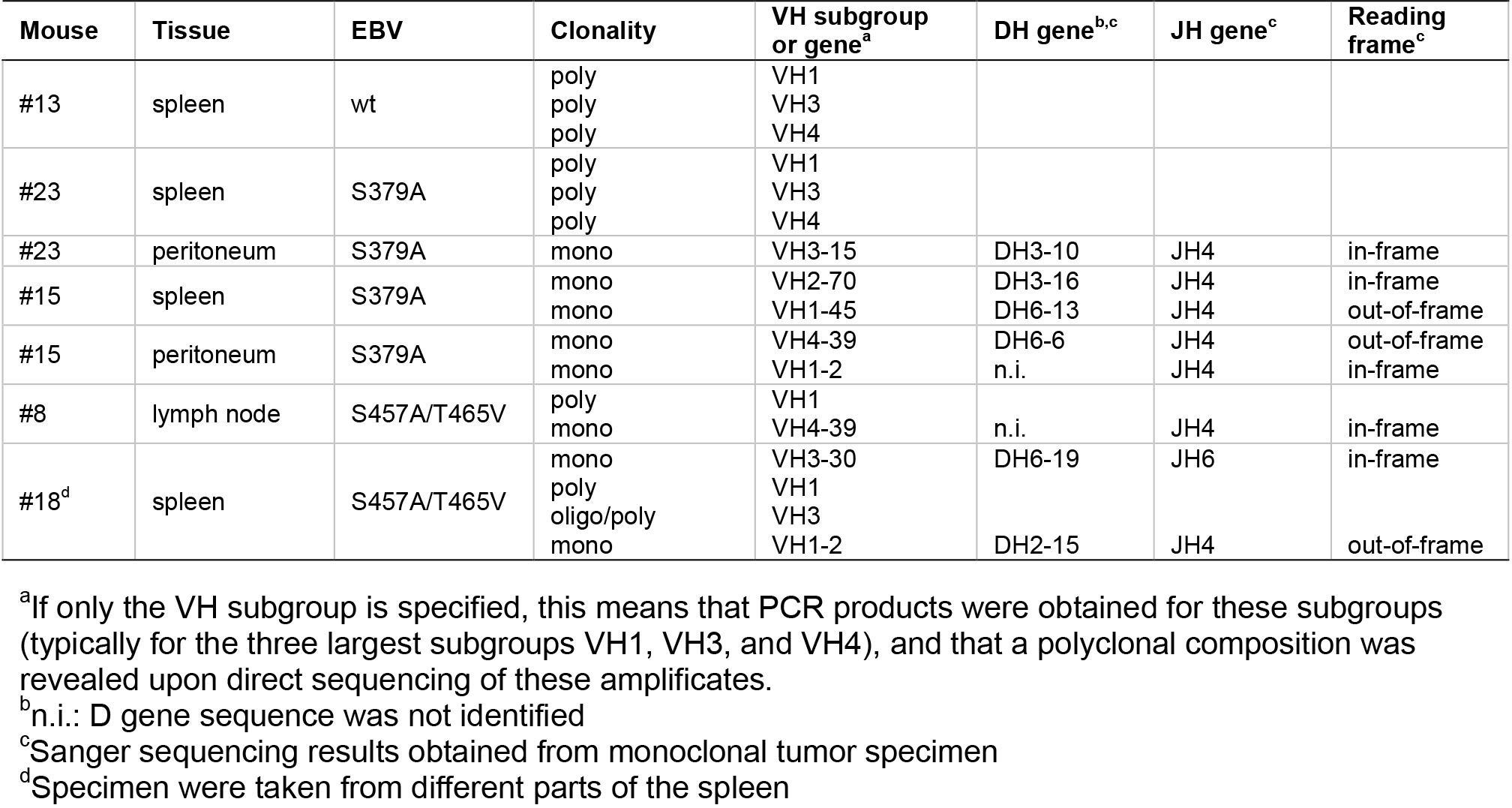
IgH V gene analysis of B cell lymphoproliferations

## DISCUSSION

### EBNA2 mutations, which impair PLK1 binding or prevent phosphorylation by PLK1, accelerate cellular proliferation and tumor formation

Our study demonstrates that PLK1 directly binds to the phosphorylated EBNA2 residue S379 and phosphorylates S457 and T465 located in the transactivation domain of EBNA2. S457A/T465V missense mutants can still bind to PLK1 but are not phosphorylated. The EBNA2 docking site S379A and EBNA2 S457A/T465V phosphorylation mutants, both show enhanced transactivation capacities with S457A/T465V being stronger than S379A. This enhanced potential might be caused in part by the elevated levels of histone acetylase and co-activator p300 that binds to the S457A/T465V mutant. Kinase active but not kinase dead PLK1 (K82M) inhibited EBNA2 wild-type activity and, to a minor extent, the S379A docking site mutant. Since the phosphorylation mutant EBNA2 S457A/T465V was not inhibited by PLK1 co-expression we conclude that EBNA2 phosphorylation, rather than sole PLK1 binding, inhibits the transactivation potential. The residual inhibition of EBNA2 S379A might be caused by PLK1 since this mutant retains some PLK1 binding activity and thus can be phosphorylated.

Both EBV mutants, which expressed EBNA2 S379A or S457A/T465T were fully immortalization competent and initiated long-term proliferating B cell cultures with accelerated cell division rates and elevated induction of the EBNA2 target gene LMP1. The impact of both mutations was further studied in humanized mice susceptible to EBV infection. Blood samples of infected mice were analyzed weekly for viral loads and immune responses of CD4^+^ and CD8^+^ T cell populations. Elevated viral loads and CD4^+^ and CD8^+^ T cell activation in the blood of EBV S457A/T465V of infected mice preceded CD8^+^ T cell expansion. On the day of sacrifice, splenic EBV loads were slightly elevated in both EBNA2 mutants compared to EBV wt infected animals. Based on earlier publications, increasing viral loads were expected to precede expansion and activation of EBV-specific CD8^+^ T cells since antigen abundance stimulates CD8^+^ T cell responses (Shultz *et al*, 2010; Strowig *et al*, 2009a). Irrespective of the early adaptive immune response to infection, more EBV mutant infected mice developed lymphomas when compared to EBV wt infections. The accelerated proliferation rates that we observed in cell culture might contribute to tumor size in mice. The enhanced transcriptional activities of the EBNA2 mutants induce higher LMP1 levels, which is a critical viral oncogene in B cell transformation and thus also might promote tumor formation in vivo. In addition, both EBNA2 mutants might also induce distinct viral and cellular RNAs, which promote tumor progression but still are to be identified.

Importantly, tumors detected in mice infected with mutant virus were predominantly monoclonal, and could be detected in the spleen as well as in the peritoneum and in lymph nodes. The elevated viral loads seen in mice infected with mutant EBV might increase tumor incidence by amplifying the number of infected B cells in these animals and thereby might initiate the lymphoproliferative disease at multiple sites but they do not explain disease progression to monoclonal B cell lymphomas. It will be interesting to study if lymphomas in lymph nodes or peritoneum are metastatic descendants of splenic tumors. The genetic or epigenetic processes driving the clonal evolution of single cells and giving rise to monoclonal tumors still need to be explored.

### PLK1 controls the growth transformation potential of EBNA2 to establish tumor free survival of latently infected hosts

Our study suggests that EBNA2 activity needs to be controlled by PLK1 in order to reduce the risk of tumor formation while latency is established in the infected animals. Since PLK1 activity peaks in mitosis, PLK1 might control EBNA2 activities preferentially during this time window. Unfortunately, since during mitosis the global cellular transcription of the condensed chromosomes and the viral episome is silenced, EBNA2 activity cannot be reliably tested in mitotic cells. There appears to be a strict biological necessity to dynamically control EBNA2 activity by multiple viral and cellular factors. Transient transfections and reporter gene studies have suggested that Cyclin B/ CDK1 or the viral lytic kinase BGLF4 can control EBNA2 activity, but these studies have not linked their findings to PLK1 activity (Yue *et al*, 2004; Yue *et al*, 2005; Yue *et al*, 2006).

PLK1 is a key control element of multiple cell cycle stages including G2/M transition, M-phase progression, and cytokinesis. In addition, it has been shown before that PLK1 can affect the activity of cellular transcription factors through several mechanisms. Phosphorylation of the tumor suppressor p53 and the related p73 protein impair the transactivation activity of both transcription factors. In p73, the substrate site of PLK1 has been mapped to the TAD (Ando *et al*, 2004; de Carcer, 2019; Koida *et al*, 2008; Martin & Strebhardt, 2006). Phosphorylation of the transcription factor FOXO1 by PLK1 causes its nuclear exclusion and thereby prevents its action (Yuan *et al*, 2014). On the other hand, PLK1 phosphorylation of the transcription factor FoxM1, a critical transactivator of mitotic gene expression, induces FoxM1 activity (Marceau *et al*, 2019). It is tempting to speculate that PLK1 might contribute to the general mitotic transcriptional shut down by inhibiting distinct transcription factor activities. In parallel, PLK1 might guide the transcriptional activity of the cell to processes relevant to mitotic progression and maintenance of genomic integrity.

In healthy immunocompetent hosts, EBNA2 is expressed in a short time window immediately post-infection before either EBNA2 expression is silenced or the EBNA2 expressing cell is eliminated by the immune system. Immunodeficient patients, suffering from profound T cell suppression, can develop aggressive EBNA2-positive B cell lymphoproliferative diseases and EBNA2 is a driving force for these tumor entities. The “primary goal” of latent viruses like EBV is to establish a balanced equilibrium of its latent and lytic phase while tumor development is an accident in its life cycle with no benefit for virus dissemination (Shannon-Lowe & Rickinson, 2019). EBNA2 obeys the intracellular signaling cues that silence its activity during a short time window of the cell cycle in order to establish latency and maintain the viral life cycle rather than promoting tumorigenesis. It is well established that high-level expression of PLK1 promotes carcinogenesis in multiple tissues (Liu *et al.*, 2017; Strebhardt, 2010; Strebhardt & Ullrich, 2006). PLK1 is also considered as an oncotarget for aggressive B cell lymphomas since it stabilizes MYC (Ren *et al*, 2018). Currently, clinical trials evaluate the safety and efficacy of PLK1 inhibitors for patient treatment. However, in some tumor types high-level PLK1 expression can suppress cancerogenesis (de Carcer, 2019; Raab *et al*, 2018).

Here we show that PLK1 is an important cellular control factor that restrains the proliferation and transformation of latently infected B cells driven by a growth program that depends on EBNA2. Since two distinct EBNA2 mutants that both target independent PLK1 related functions of the EBNA2/PLK1 complex promote cancerogenesis, we conclude that PLK1 might act as a tumor suppressor in EBNA2 driven pathogenesis. Based on our results, the development and therapeutic use of PLK1 inhibitors should be re-considered and closely monitored with respect to potential adverse effects in the context of the prevalent latent EBV infections in the population.

## MATERIALS AND METHODS

### Cell culture

EBV infected long-term cultures (LCLs), DG75 cells (Ben-Bassat *et al*, 1977), DG75^Dox HA-EBNA2^ (Glaser *et al.*, 2017), Raji (Pulvertaft, 1964), and HEK293 cells were cultured in RPMI 1640 supplemented with 10% fetal bovine serum (FBS), 1% non-essential amino acids, 2 mM L-glutamine, 1 mM sodium pyruvate, 100 U/ml penicillin, 100 μg/ml streptomycin, 100 nM sodium selenite and 50μM α-thioglycerols at 37°C in 6% CO_2_ atmosphere. Media for HEK293 cells transfected with recombinant EBV and DG75^Dox HA-EBNA2^ contained 1 μg/ml puromycin.

### Purification of human primary B cell and LCL establishment

Human primary B cells were isolated from adenoid from routine adenoidectomy were obtained from the Department of Otorhinolaryngology, Klinikum Grosshadern, Ludwig Maximilians University of Munich, and Dritter Orden Clinic, Munich-Nymphenburg, Germany. All clinical samples were made fully anonymous. To isolate human primary B cells, T cells were depleted by erythrocyte rosetting using sheep red blood cells and B cells were separated by Ficoll density gradient centrifugation as recommended by the manufacturer (GE Healthcare). The remaining erythrocytes were lysed in red blood cell lysis buffer (155 mM NH_4_Cl, 10 mM KHCO_3_, 0.1 mM EDTA). Cells were co-stained by anti-CD3^+^ (UCHT1; BD Pharmingen) and anti-CD19^+^(HIB19; BD Pharmingen) antibodies and analyzed by flow cytometry.

To generate LCLs, primary human B cells were infected with recombinant EBV at a ratio of 1 green raji unit (GRU) to 10 cells for 48 h and cultivated in medium containing 0.5 μg/ml cyclosporine A for 2 weeks before routine cell culture conditions were applied.

### Antibodies, western blot, immunoprecipitation, and GST-pull down

Monoclonal antibodies specific for His_6_ (2F12), EBNA2 (R3), Glutathione S transferase (6G9), HA-tag (3F10), BSA (3C5), LMP1 (1G6) were provided by the antibody facility of the Helmholtz Center Munich. Commercial providers were: GAPDH (Mab374; Merck Millipore), Flag (M2; Sigma Aldrich), PLK1 (ab17056; Abcam), GFP (7.1 and 13.1; Roche) and a p300 specific serum (C20; Santa Cruz). For immunoprecipitations and Western blotting the equivalent of 1×10^7^ cells was lysed in 500 μl of lysis buffer (1% NP40, 10 mM Tris-HCl pH 7.4, 3% glycerol, 150 mM NaCl, 1 mM EDTA) supplemented with cOmplete protease inhibitor and PhoStop phosphatase inhibitor (Roche) incubated for 30 min at 4°C with constant rolling and for 30 min on ice. The lysate was cleared by centrifugation (16000 g, 15 min). 1 μg of purified antibodies or 100 μl of hybridoma supernatants were coupled to Protein A or G beads and added to the cleared lysates, incubated for 2 hours at 4°C, washed with lysis buffer and the protein was eluted with Laemmeli buffer. For GST pulldown, antibody coated beads were replaced by GST fusion protein coated beads. 20 μg protein of total cell lysates or 5×10^6^ cell equivalents of one immunoprecipitation were loaded per lane. Signals on Western blots were detected by enhanced chemiluminescence (GE Healthcare) and exposed to films or Fusion FX (Vilber Lourmat).

### Construction of plasmids

All the plasmid used in the study were cloned based on conventional PCR, restriction digestion, and ligation. Mutated alleles were generated by overlap PCR adapted from the previous protocol (reviewed in Francis *et al*, 2017). In essence of overlap PCR is based on four strategically designed primers. Internally positioned primers must contain complementary sequences to each other, and both of them must contain the mutation of interest, like a substitution, a deletion, or an insertion. The flanking primers might contain restriction enzyme recognition sites to facilitate the cloning of the amplified fragment. Two steps were performed, in the first round of PCR reactions using the forward primer of the flanking primers with the reverse primer of the internally positioned primers and vice versa, respectively. The resulting amplified short fragments worked as templates when mixed with the flanking primer pairs, which results in amplification of the final long fragment with the desired mutation in the second round of PCR. The second PCR product is digested and inserted into a corresponding vector. For each plasmid used in the study detail information about primer pairs, template, and vector is shown in Table S1.

### Expression and purification of His- or GST-tagged proteins

His6-tagged and GST-tagged proteins were expressed in E. coli Rosetta (DE3) cells and purified according to manufacturer’s instructions using Ni-chelate agarose (Quiagen) or glutathione coupled Sepharose 4B beads (GE Healthcare).

### Kinase assay in vitro

Purified protein or protein coupled to beads were incubated for 30 min at 37°C with recombinant active Cyclin B/CDK1 (100 ng) or PLK1 (50 ng) in the presence of 1mM normal ATP or plus 0.25 mCi/ml γ-^32^P labeled ATP in PK buffer (50 mM Tris-HCl, 10 mM MgCl_2_, 0.1 mM EDTA, 2 mM DTT, 0.01% Brij 35, pH 7.5) in a total reaction volume of 20 μl.

### EBV BAC recombineering

All recombinant EBV strains used in this study were generated by a two-step selection protocol using the λ-prophage-based heat inducible Red recombination system expressed in E. coli strain SW105 (Pich *et al.*, 2019; Wang *et al.*, 2009; Warming *et al*, 2005). For the first step, a Kan/rpsL expression cassette was flanked by 50 nt EBV sequences of the respective EBNA2 gene locus by PCR using the template p6012. The resulting PCR product was used to insert the Kan/rpsL cassette by homologous recombination into the specific EBV/EBNA2 target site by transformation and kanamycin (30 μg/ml) and chloramphenicol (12.5 μg/ml) selection of SW105 pre-transformed with the recombinant target EBV plasmid. As a second step, a synthetic DNA fragment or PCR product carrying the desired mutation flanked by ~300 nt of the genomic viral sequence was used to replace the Kan/rpsL cassette by homologous recombination to generate the final mutant EBV plasmid by streptomycin (1 mg/ml) and chloramphenicol (12.5 μg/ml) selection. For each BACmid used in the study, detailed information about primer pairs, template, and vector are shown in Table S1.

### Production of recombinant virus

HEK293 transfectants carrying the recombinant virus plasmid were induced for virus production by co-transfection of 0.5 μg of the plasmids p509 encoding BZLF1 and p2670 encoding BALF4 per one 6-well in 3 ml cell cultures. The supernatants of the transfectants were harvested 3 days post-transfection and passed through a 0.8 μm filter. For quantification of viral titers, 1×10^5^ Raji cells were infected with serial dilutions of viral supernatants in 2 ml cultures and the percentage of GFP positive cells was determined by FACS analysis 3 days post-infection. The concentration of viral stocks was expressed as the number of green Raji units (GRU).

### Isothermal titration calorimetry

Experiments were performed on a ITC200 instrument in triplicates and analyzed with the Malvern software. 100 μM PBD was provided in the cell and titrated with 1 mM concentration of wild type ((PNTSSPS) or phosphopeptide (PNTSpSPS) with 25 times 1.5 μL injections at 25°C.

### Generation and infection of humanized mice

NOD scid γ_c_^−/−^ (NSG) mice obtained from the Jackson Laboratories were bred and maintained under specific pathogen-free conditions at the Institute of Experimental Immunology, University of Zurich. CD34^+^ human hematopoietic progenitor cells were isolated from human fetal liver tissue (obtained from Advanced Bioscience Resources) using the CD34 MicroBead Kit (Miltenyi Biotec) following the protocol provided by the manufacturer. Newborn NSG mice (age: 1 to 5 days) were irradiated with 1 Gy by use of an x-ray source. 1 - 3 × 10^5^ CD34+ human hematopoietic progenitor cells were injected intrahepatically 5 to 7 hours after irradiation. Reconstitution of mice with human immune system components was investigated 10 - 12 weeks after engraftment by flow cytometry for the cell surface expression of huCD45 (BV605 or Pacific Blue, clone HI30; Biolegend), huCD3 (PE, clone UCHT1; BV785, clone OKT3; Biolegend), huCD19 (PE-Cy7, clone HIB19; Biolegend) and (PE-Texas Red, Clone SJ25-C1), huCD4 (APC-Cy7, clone RPA-T4; Biolegend), huCD8 (PerCP, clone SK1; Biolegend), huNKp46 (APC, clone 9E2; BD) and HLA-DR (FITC or PE-Cy7 clone L243; Biolegend) on PBMCs. 12 - 16 weeks after engraftment, humanized mice were infected intraperitoneally with 1×10^5^ GRU of EBV wt, EBV S379A or EBV S457A/T465V. For each experiment, a different cohort of mice reconstituted with CD34^+^ cells derived from one donor was generated. The animals were ascribed to a distinct experimental group ensuring similar ratios of males to females and similar reconstitution levels and immune cell activation in the peripheral blood. 5 weeks after infection mice were sacrificed if not necessitated earlier by the regulations of our experimental animal license as a consequence of general health conditions or weight loss over 20%. For analysis of the experiments, only those mice that showed two of the following signs of infection were regarded as infected and included in the analysis: (i) Viral loads in spleen, (ii) viral loads in blood at one time-point during the experiment, (iii) EBNA2^+^ cells in spleen as evaluated by histology. The respective animal protocol (ZH159-17/ZH008-20) was approved by the veterinary office of the canton of Zurich, Switzerland.

### Whole blood and spleen preparations for immune phenotyping

Whole blood of mice was collected from the tail vein and prepared for immunophenotyping by lysing erythrocytes with NH_4_Cl. Spleens of mice were mashed, subsequently filtered with a 70 μm cell strainer, and afterwards mononuclear cells were separated using Ficoll-Paque gradients. Total cell counts were determined from purified mononuclear cell suspensions using a DxH500 Hematology Analyzer (Beckman Coulter). Purified cell suspensions were stained for 30 - 40 minutes at 4°C in the dark with a master mix of the respective antibodies followed by a washing step in PBS. The stained cells were analyzed in an LSR Fortessa cytometer (BD Biosciences). Flow cytometry data were analyzed using the FlowJo software.

### EBV DNA Quantification in Tissue

Total DNA from splenic tissue and whole blood was extracted using DNeasy Blood & Tissue Kit (QIAGEN) and NucliSENS (bioMérieux), respectively, according to manufacturer’s instructions. Quantitative analysis of EBV DNA in humanized mouse spleens and blood was performed by a TaqMan (Applied Biosystems) real-time PCR as described previously (Berger et al., 2001) with modified primers (5’-CTTCTCAGTCCAGCGCGTTT-3’ and 5’-CAGTGGTCCCCCTCCCTAGA-3’) and the fluorogenic probe (5’-(FAM)-CGTAAGCCAGACAGCAGCCAATTGTCAG-(TAMRA)-3’) for the amplification of a 70-base pair sequence in the conserved BamHI W fragment of EBV. Real-time PCR was run on a ViiA 7 Realtime PCR system or a CFX384 Touch Real-Time PCR Detection System and samples were analyzed in duplicates.

### IgV gene rearrangement sequence analysis

For PCR analysis of rearranged immunoglobulin genes, macroscopically visible tumors were dissected and cryofixated in Tissue Tek^®^ O.C.T.™ (VWR, Cat# SAKU4583) on dry ice.

DNA was isolated from 10-20 8-μm frozen tissue sections of biopsies from humanized mice using the PureGene DNA isolation kit (Qiagen, Hilden, Germany). PCR was performed with six framework region 1 subgroup specific primers and a JH primer mix (3’ JH mix) for 35 cycles in six separate reactions, using 300 ng of DNA per reaction. Primer sequences have been published before (Kuppers *et al*, 2019). PCR products were purified from agarose gels and Sanger sequenced with the IGHV primers used for PCR and the BigDye Sequencing Kit (ABI, Heidelberg, Germany) on an Applied Biosystems 3130 Genetic Analyzer (ABI). Sequences were evaluated with the IMGT/V-Quest software (http://www.imgt.org/IMGT_vquest/input).

### Mass Spectrometry of EBNA2 co-immunoprecipitates

DG75 cells were transfected with a C-terminal HA-tagged EBNA2 (plasmid: pAG155) using electroporation. After 48 h, cells were lysed in NP-40 lysis buffer (1% NP40, 10 mM Tris-HCl pH 7.4, 3% glycerol, 150 mM NaCl, 1 mM EDTA supplemental with cOmplete protease and PhoStop phosphatase inhibitor). Cell debris was depleted by centrifugation. The cell extract was incubated with protein G beads covalently coupled with an HA-specific antibody. Immunoprecipitates were eluted with Lämmeli buffer.

After reduction and alkylation using DTT and IAA, the proteins were centrifuged on a 30 kDa cutoff filter device, washed thrice with UA buffer (8 M urea in 0.1 M Tris/HCl pH 8.5) and twice with 50 mM ammonium bicarbonate. The proteins were digested for 16 h at 37°C using 1 μg trypsin. After centrifugation (10 min at 14,000× g), the eluted peptides were acidified with 0.5% Trifluoroacetic acid and stored at −20°C.

LC-MS/MS analysis was performed on a LTQ Orbitrap XL mass spectrometer (Thermo Scientific, Waltheim, MA, USA) online coupled to an Ultimate 3000 nano-RSLC (Thermo Scientific).Tryptic peptides were automatically loaded on a C18 trap column (300 μm inner diameter (ID) × 5 mm, Acclaim PepMap100 C18, 5 μm, 100 Å, LC Packings) prior to C18 reversed phase chromatography on the analytical column (Acclaim PepMap C18, 50 μm ID × 250 mm, 2 μm, 100 Å, LC Packings) at 300 nL/min flow rate in a 140 min acetonitrile gradient from 4 to 30% in 0.1% formic acid. Profile precursor spectra from 300 to 1500 m/z were recorded in the orbitrap with a maximum injection time of 500 ms. TOP10 fragment spectra of charges 2 to 7 were recorded in the linear ion trap with a maximum injection time of 100 ms, an isolation window of 2.0 m/z, a normalized collision energy of 35 and a dynamic exclusion of 60 s. Raw files were analyzed using Progenesis QI for proteomics (version 2.0, Nonlinear Dynamics, part of Waters). Features of charges 2–7 were used and all MS/MS spectra were exported as mgf file. Peptide searches were performed using Mascot search engine (version 2.5.1) against the Ensembl Human protein database (100158 sequences, 37824871 residues). Search settings were: 10 ppm precursor tolerance, 0.6 Da fragment tolerance, one missed cleavage allowed. Carbamidomethyl on cysteine was set as a fixed modification, deamidation of glutamine and asparagine allowed as variable modification, as well as oxidation of methionine. Applying the percolator algorithm with a cut-off of 13 and p < 0.05 resulted in a peptide false discovery rate (FDR) of 1.54%. Search results were reimported in the Progenesis QI software. Proteins were quantified by summing up the abundances of all unique peptides per protein. Resulting protein abundances were used for the calculation of fold-changes and statistics values.

### Mass Spectrometry on purified EBNA2 phosphorylated by PLK1 in vitro

EBNA2 and GST-fused EBNA2 proteins, purified from bacteria, were subjected to PLK1 kinase assays. 50 μg EBNA2 or 130 μg GST-fused EBNA2 protein were incubate in 1x PK buffer (NEB, 50 mM Tris-HCl, 10 mM MgCl_2_, 0.1 mM EDTA, 2 mM DTT, 0.01% Brij 35, pH 7.5), 1.25 mM ATP in presence or absence of 250 ng active PLK1 (SignalChem) at 37°C for 1 h. All reactions were carried out in a total volume of 20 μL and then were quenched with 5 μL 5x Lämmli sample buffer.

Proteins were separated by SDS-PAGE, and stained with Coomassie colloidal blue. Bands corresponding to EBNA2 full length (86 kD) or GST-EBNA2 C-terminal fragment (28 kD) were sliced out from the gel lane, and proteins were then reduced, alkylated, and digested with either trypsin or GluC (Roche), as previously described (Shevchenko *et al*, 1996).

Dried peptides were reconstituted in 0.1% FA/2% ACN and subjected to MS analysis using a Dionex Ultimate 3000 UHPLC+ system coupled to a Fusion Lumos Tribrid mass spectrometer (Thermo Fisher). Peptides were delivered to a trap column (75 μm × 2 cm, packed in-house with 5 μm Reprosil C18 resin; Dr. Maisch) and washed using 0.1% FA at a flow rate of 5 μL/min for 10 min. Subsequently, peptides were transferred to an analytical column (75 μm × 45 cm, packed in-house with 3 μm Reprosil C18 resin, Dr. Maisch) applying a flow rate of 300 nL/min. Peptides were chromatographically separated using a 50 min linear gradient from 4% to 32% solvent B (0.1% FA, 5% DMSO in ACN) in solvent A (0.1% FA in 5% DMSO). The mass spectrometer was operated in data-dependent mode, automatically switching between MS and MS/MS. Full-scan MS spectra (from *m/z* 360 to 1500) were acquired in the Orbitrap with a resolution of 60,000 at *m/z* 200, using an automatic gain control (AGC) target value of 5e5 charges and maximum injection time (maxIT) of 10 ms. The 10 most intense ions within the survey scan were selected for HCD fragmentation with normalized collision energy set to 28%.

Isolation window was set to 1.7 Th, and MS/MS spectra were acquired in the Orbitrap with a resolution of 15,000 at *m/z* 200, using an AGC target value of 2e5, and a maxIT of 75 ms. Dynamic exclusion was set to 20 s.

Peptide and protein identification was performed using MaxQuant (version 1.5.3.30) with its built in search engine Andromeda (Cox & Mann, 2008). Spectra were searched against a SwissProt database, either the *Spodoptera frugiperda* (OX 7108 - 26,502 sequences) or *Escherichia coli* (UP000002032 – 4,156 sequences), supplemented with the EBNA2 protein sequence. Enzyme specificity was set to Trypsin/P or GluC accordingly, and the search included cysteine carbamidomethylation as a fixed modification, protein N-term acetylation, oxidation of methionine, and phosphorylation of serine, threonine, tyrosine residue (STY) as variable modifications. Up to two and three missed cleavage sites were allowed for trypsin and GluC, respectively. Precursor tolerance was set to 4.5 ppm (after MS1 feature re-calibration), and fragment ion tolerance to 20 ppm. The match between runs feature was enable. Peptides identification were further filtered for a minimum Andromeda score of 20 or 40, for unmodified and modified (phosphorylated) sequences, respectively. A site localization probability of at least 0.75 was used as the threshold for confident localization (Vizcaino *et al*, 2013; Vizcaino *et al*, 2016).

### Cell proliferation assay

Human adenoid primary B cells were stained with CellTrace Violet according to the manufacturer’s instructions (Thermo Fisher Scientific). Proliferation of CD19^+^ B cells was monitored by flow cytometry using BD Fortessa and the data were analyzed using the FlowJo software (Version 10.5.3).

### Dual luciferase assay

5x 10^6^ DG75 cells were electroporated with 1.5 μg EBNA2 expression plasmids and the luciferase construct 1.5 μg pGa981.6 (Minoguchi *et al.*, 1997) carrying a multimerized CBF1 binding site to measure EBNA2 activity and 0.2 μg renilla luciferase expression plasmid. 24 h post electroporation, cells were washed, pelleted and lysed in 100 μL lysis buffer for 30 min on ice. Cell extracts were tested by the dual luciferase assay according to the manufacturer’s instructions (Promega).

## RESOURCES

### Data Availability

Phosphorylation of EBNA2 by PLK1: The mass spectrometry proteomics data have been deposited in the ProteomeXchange Consortium via the PRIDE partner repository (Link: https://www.ebi.ac.uk/pride/login - Username: - Password: wZd0Qfaz) with the dataset identifier PXD022970 (Vizcaino *et al.*, 2013; Vizcaino *et al.*, 2016).

## ACKNOWLEDGEMENT

We thank Dagmar Pich, Yen-Fu Adam Chen, Ezgi Akidil for all the excellent advice they gave.

We thank Kerstin Heise, Michaela Kroetz-Fahning and Andreas Klaus for expert technical assistance. This project was supported by the Wilhelm Sander-Stiftung (Grant 2015.165.1) to Bettina Kempkes. Xiang Zhang is supported by the China Scholarship Council (CSC No.: 201603250052). Cristian Münz is financially supported by Cancer Research Switzerland (KFS-4091-02-2017 and KFS-4962-02-2020), KFSP-PrecisionMS and HMZ ImmunoTargET of the University of Zurich, the Cancer Research Center Zurich, the Vontobel Foundation, the Baugarten Foundation, the Sobek Foundation, the Swiss Vaccine Research Institute, Roche, Novartis and the Swiss National Science Foundation (310030B_182827, 310030L_197952/1 and CRSII5_180323)

## AUTHOR CONTRIBUTIONS

Conceptualization: XZ, MS, CM, RK, BKe; Formal analysis and data curation: AMo, PG, SMH, Methodology: XZ, PS, MR, KS, CM; Funding acquisition: BKe, CM; Investigation: XZ, PS, AMo, PG, AMu, ST, CKR, SB, RK; Resources: WH, MR, MS, CM, KS, BKu, CM; Supervision: WH, RK, MS, CM, BKe; Visualization: XZ, PS, AMo; BKe; Writing-original draft: XZ, BKe; Writing-review&editing: XZ, MS, CM, PG, PS, AMo, BKe; RK;

---

**Table EV1.** 19 candidate EBNA2 associated proteins identified by label free mass spectrometry

| ensemble-ID | Peptide count<sup>a</sup> | Peptides used for quantification<sup>b</sup> | Confidence score<sup>c</sup> | Anova (p) | Accumulation<sup>d</sup> | Symbol | Replicate 1<sup>e</sup><br>vector (empty) | Replicate 2<sup>e</sup><br>vector (empty) | Replicate 3<sup>e</sup><br>vector (empty) | Replicate 1<sup>e</sup><br>EBNA2 wt | Replicate 2<sup>e</sup><br>EBNA2 wt | Replicate 3<sup>e</sup><br>EBNA2 wt | Description |
| --- | --- | --- | --- | --- | --- | --- | --- | --- | --- | --- | --- | --- | --- |
| ENSP00000248958 | 1 | 1 | 48 | 0.00 | 1652 | SDF2L1 | 27 | 0 | 0 | 4799 | 33098 | 6503 | stromal cell-derived factor 2-like 1 |
| ENSP00000247020 E | 2 | 2 | 54 | 0.01 | 762 | SDF2 | 26 | 0 | 0 | 2392 | 14742 | 2515 | stromal cell-derived factor 2 |
| ENSP00000342070 | 2 | 2 | 85 | 0.04 | 177 | CTSB | 56 | 0 | 36 | 11264 | 330 | 4616 | cathepsin B |
| ENSP00000380376 | 1 | 1 | 34 | 0.01 | 164 | PAXIP1 | 13 | 27 | 2 | 338 | 4697 | 1748 | PAX interacting (with transcription-activation domain) protein 1 |
| ENSP00000311766 | 13 | 1 | 336 | 0.04 | 125 | ATAD3B | 59 | 37 | 0 | 469 | 9408 | 2205 | ATPase family, AAA domain containing 3B |
| ENSP00000300093 | 7 | 7 | 172 | 0.01 | 77 | PLK1 | 1610 | 407 | 198 | 14583 | 122709 | 32353 | polo-like kinase 1 |
| ENSP00000265028 | 5 | 5 | 207 | 0.01 | 46 | DNAJB11 | 1645 | 2063 | 2421 | 20887 | 211659 | 49942 | DnaJ (Hsp40) homolog, subfamily B, member 11 |
| P12978.1 | 5 | 5 | 162 | 0.01 | 26 | EBNA2 | 5841 | 3949 | 3629 | 41868 | 251544 | 59586 | Epstein-Barr nuclear antigen 2 |
| ENSP00000305815 | 1 | 1 | 26 | 0.01 | 26 | CBF1/RBPJ | 410 | 121 | 93 | 1602 | 10431 | 4182 | recombination signal binding protein for immunoglobulin kappa J region |
| ENSP00000261893 | 7 | 7 | 210 | 0.01 | 17 | LACTB | 5688 | 3130 | 2773 | 21048 | 129818 | 47963 | lactamase, beta |
| ENSP00000368030 | 18 | 6 | 544 | 0.03 | 11 | ATAD3A | 13872 | 9774 | 4851 | 41200 | 213392 | 45267 | ATPase family, AAA domain containing 3A |
| ENSP00000446596 | 3 | 2 | 105 | 0.05 | 10 | DYNLL1 | 13993 | 9084 | 6329 | 25470 | 217534 | 46037 | dynein, light chain, LC8-type 1 |
| ENSP00000277900 | 5 | 4 | 100 | 0.05 | 7 | ADD3 | 6551 | 765 | 819 | 12200 | 35354 | 9851 | adducin 3 (gamma) |
| ENSP00000306223 | 2 | 2 | 32 | 0.03 | 4 | SNRPN | 508 | 154 | 254 | 1595 | 1306 | 639 | small nuclear ribonucleoprotein polypeptide N |
| ENSP00000280326 | 2 | 2 | 47 | 0.05 | 2 | CCT5 | 950 | 950 | 1819 | 1867 | 3216 | 4077 | chaperonin containing TCP1, subunit 5 (epsilon) |
| ENSP00000295688 | 2 | 2 | 45 | 0.03 | 2 | CCT3 | 1269 | 1133 | 1066 | 1579 | 1979 | 2715 | chaperonin containing TCP1, subunit 3 (gamma) |
| ENSP00000261182 | 1 | 1 | 35 | 0.01 | 2 | NAP1L1 | 1081 | 843 | 707 | 1562 | 1580 | 1540 | nucleosome assembly protein 1-like 1 |
| ENSP00000272163 | 1 | 1 | 29 | 0.01 | 2 | LBR | 1550 | 1597 | 1653 | 2133 | 3232 | 2798 | lamin B receptor |
| ENSP00000238081 | 4 | 3 | 123 | 0.00 | 2 | YWHAQ | 5744 | 5869 | 7103 | 9673 | 10151 | 10026 | tyrosine 3-monooxygenase/tryptophan 5-monooxygenase activation protein, theta polypeptide |

**Figure EV1.**
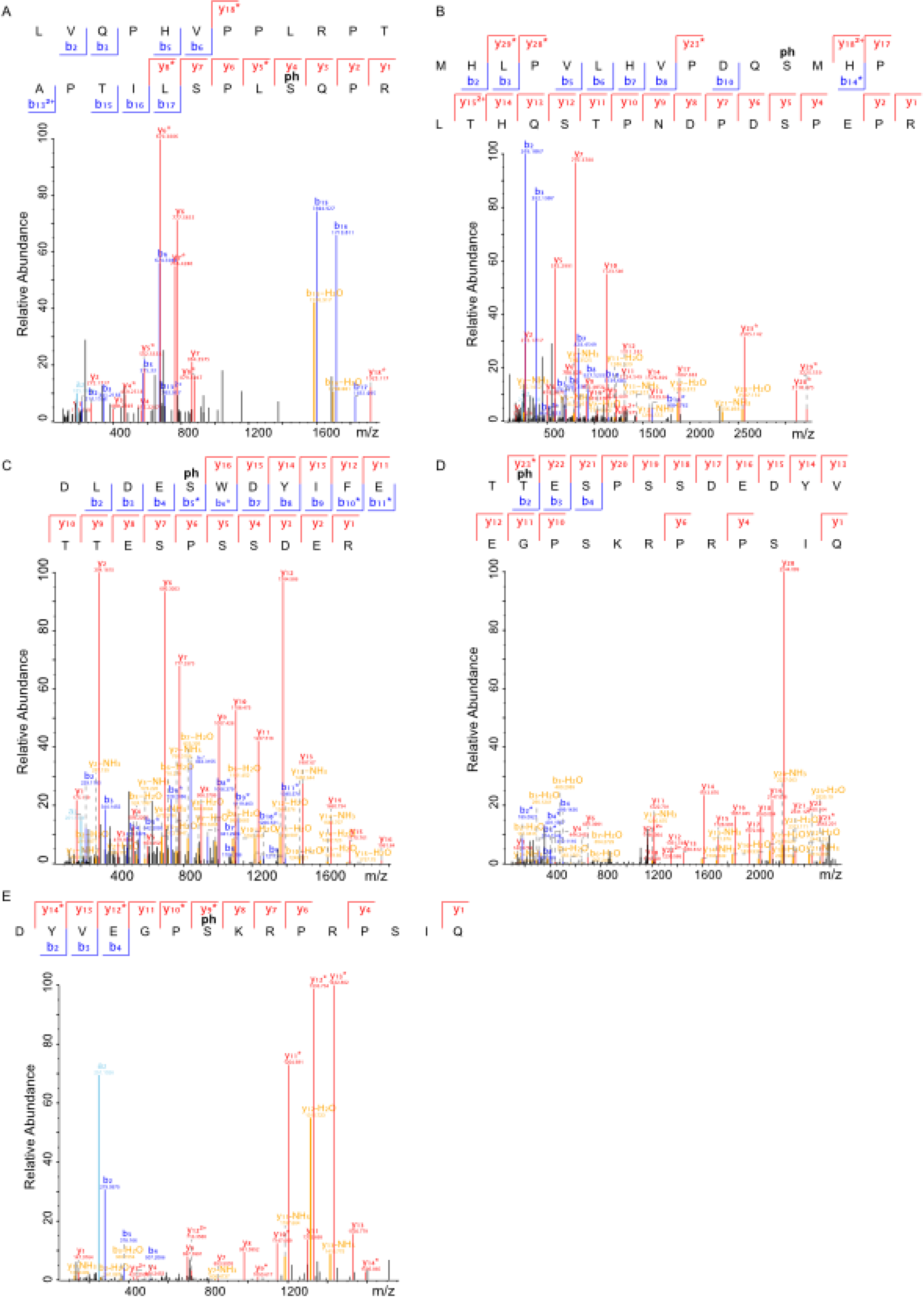
Annotated HCD MS/MS spectra of phosphopeptides. (A) LVQPHVPPLRPTAPTILSPLSQPR, (B) MHLPVLHVPDQSMHPLTHQSTPNDPDSPEPR, (C) DLDESWDYIFETTESPSSDER, (D) TTESPSSDEDYVEGPSKRPRPSIQ, and (E) DYVEGPSKRPRPSIQ, bearing 5 confidentially localized phosphorylation sites, S184, 258, 457, T465, and S479, respectively. The “ph” denotes phosphosites localized. The a-, b-, and y-ions are in pale blue, dark blue, and red, respectively. Ions with neutral losses are in orange, internal fragment ions in purple, ammonium ion in green, and side-chain loss in turquoise. The asterisk (*) denotes loss of H_3_O_4_P with a delta mass of 97.9768 from the phosphorylated fragment ion.

**Figure EV2.**
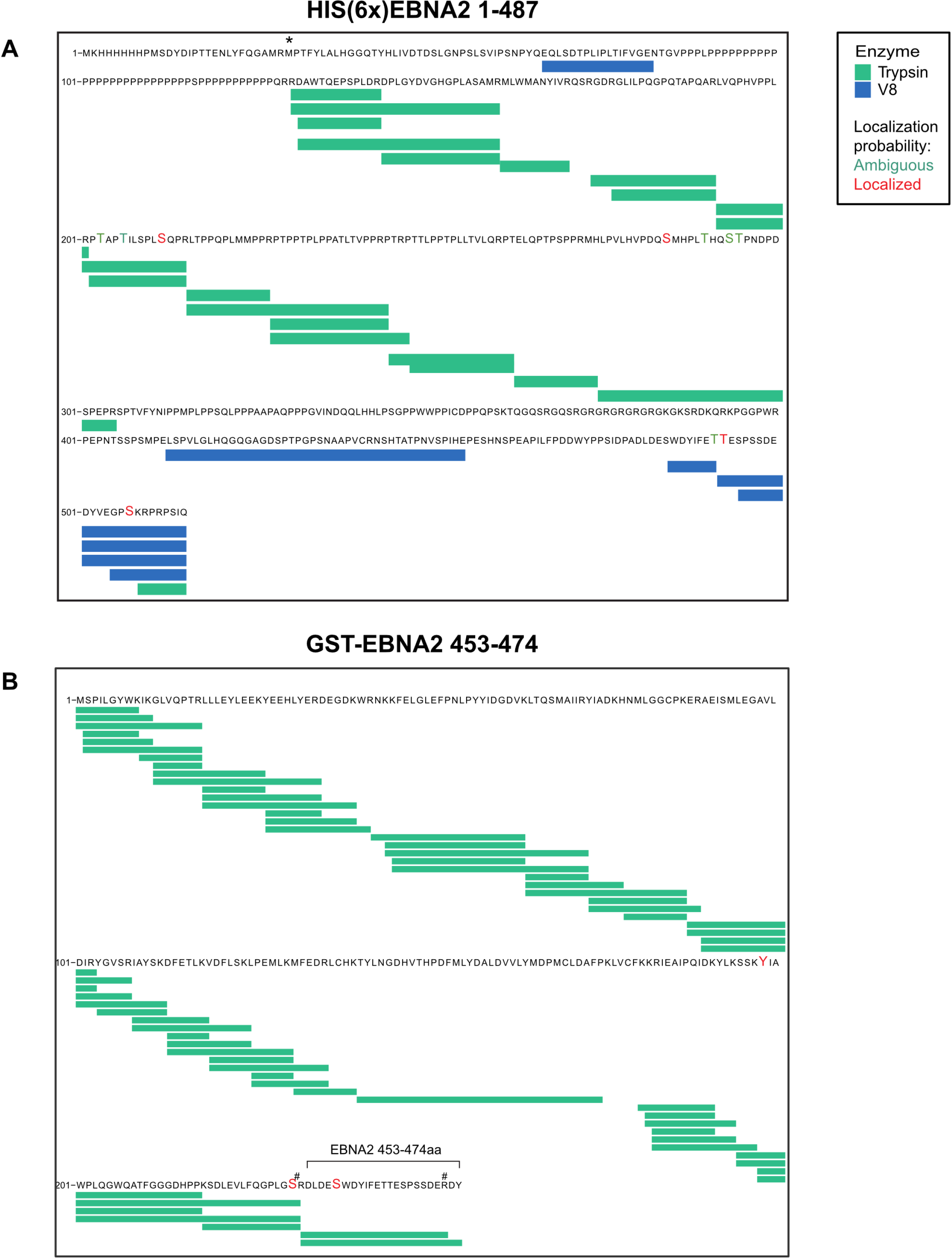
Overview of sequence coverage and phosphorylation sites identified by mass spectrometry. (A) 6x His-tagged EBNA2 (A) and (B) GST-EBNA2 453-474 phosphorylated by PLK1. Proteins were expressed in E. coli and extracted after SDS-PAGE separation, digested by trypsin (green bars) and V8 (blue bars) in parallel and submitted to LC-MS/MS (enlarged letters in red or green). The sign # denotes Arg (R) inserted to facilitate fragmentation. The asterisk (*) denotes the initial Met (M) of EBNA2. Proteins before phosphorylation by PLK1 were studied in parallel but no phosphorylation sites were identified. The sequence corresponds to UniProt ID: P12978.1.

**Figure EV3.**
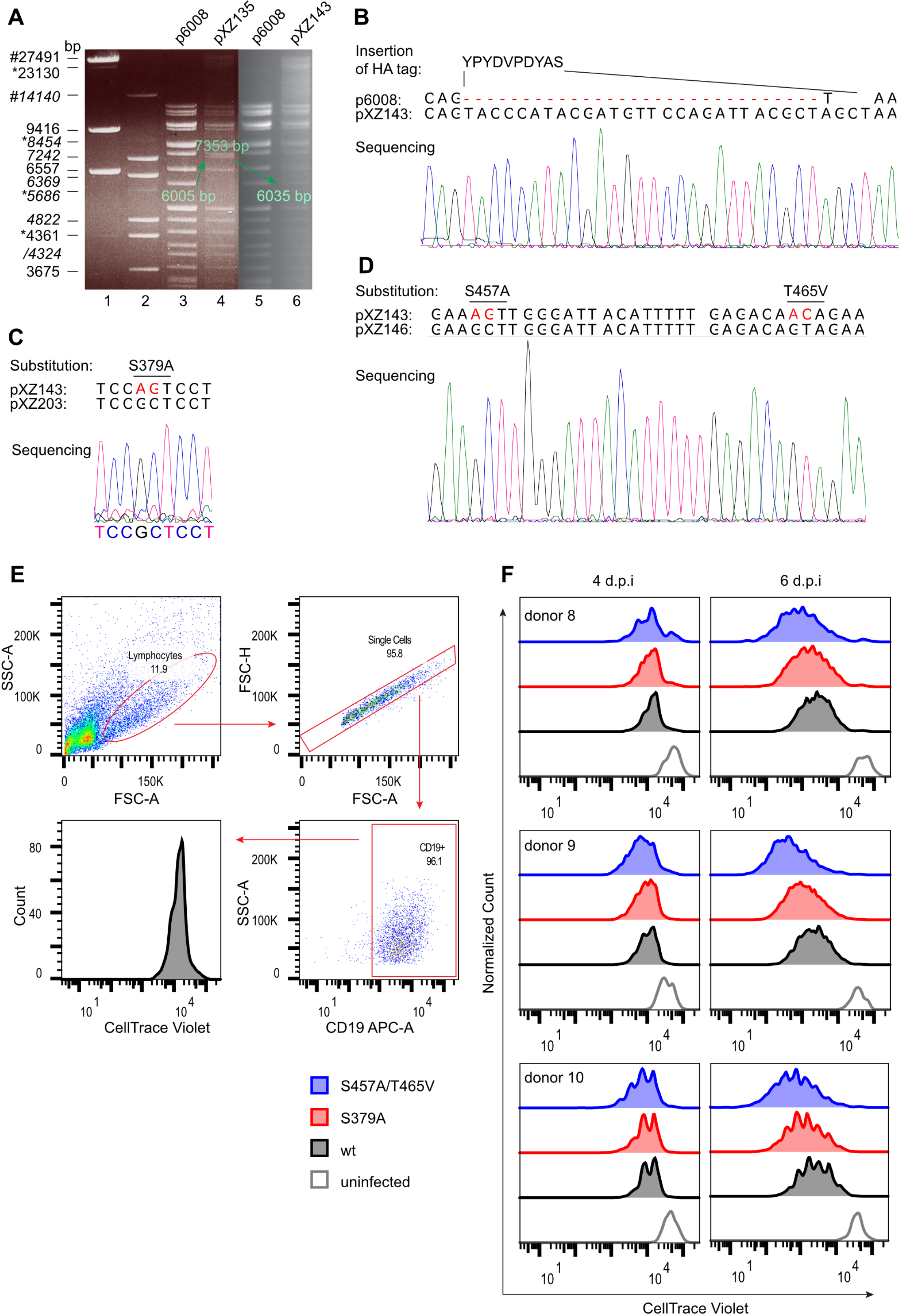
Construction of EBV BACmids carrying HA-tagged EBNA2 mutants impaired for PLK1 binding or PLK1 phosphorylation and functional tests in cell culture. (A) Electrophoretic separation of the restriction digest of EBV Bac DNA: p6008 (precursor), pXZ135 (insertion of Kan/rpsL as a precursor for EBV HA-EBNA2) and pXZ143 (EBV HA-EBNA2=EBV wt). The arrows highlight distinct fragments that characterize the individual BACs in size upon Kan/rpsL insertion and deletion (6005 bp → 7353 bp → 6035 bp). Molecular markers: λ DNA-Hind III digest (nonitalics) and λ DNA-BstE II digest (italics). (#) denotes the fragment derived from fragments denoted by asterisks (*). (B) Sanger sequencing of pXZ143 to confirm the insertion of the HA-tag into EBNA2 in the backbone of p6008 (C, D) Sanger sequencing of pXZ203 (C) and pXZ146 (D) to confirm the substitution of S379A and S457A/T465V, respectively. (E) Gating strategy of cell trace violet stained active B cells after EBV infection. (F) Adenoid B cells were stained with cell trace violet before they were infected with EBV mutants as indicated and analyzed by flow cytometry.

**Figure EV4.**
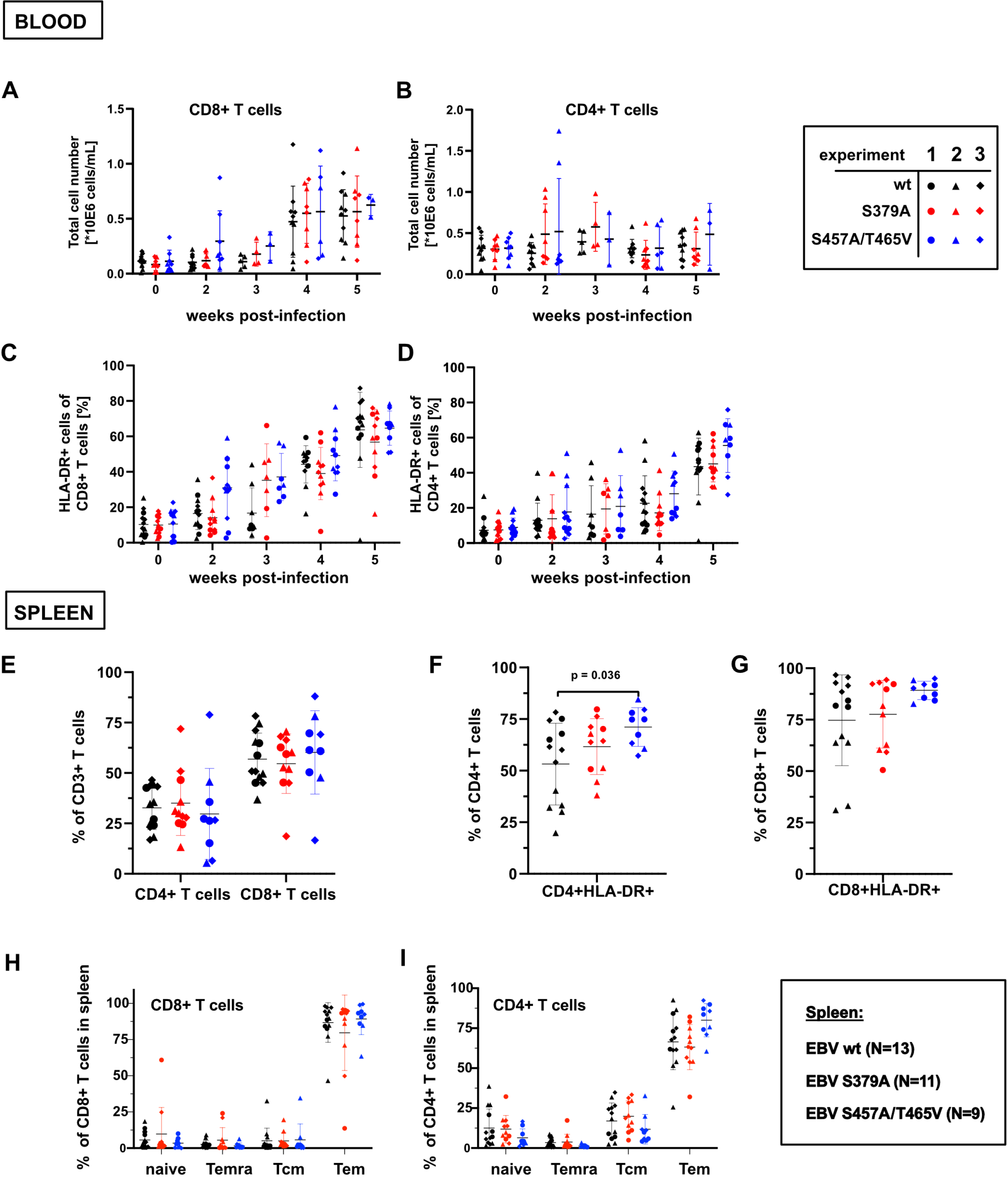
T cell subpopulations of blood and spleen of infected mice. Flow-cytometric analyses of (A) CD8^+^ or (B) CD4^+^ T cell percentages and (C) CD8^+^ and (D) CD4^+^ T cell activation in the blood of mice infected with wt or mutant EBV over a period of five weeks. (E) Percentages of CD4^+^ and CD8^+^ subpopulations and activation of (F) CD4^+^ and (G) CD8^+^ subpopulations. Analyses of the spleens for (H) CD8^+^ and (I) CD4^+^ naïve, terminally differentiated (Temra), central memory (TCM) and effector memory (Tem) memory T cell subpopulations. (The shape of data points indicates to which cohort (experiment 1, 2 or 3) the respective animal belongs while the color indicates the EBV strain. Statistical significance was tested using the Mann-Whitney U Test with Holm-Sidak correction for multiple comparisons.

## APPENDIX

Supplementary Table S1 (List of oligonucleotides used in this study)

## REFERENCES

Ando K, Ozaki T, Yamamoto H, Furuya K, Hosoda M, Hayashi S, Fukuzawa M, Nakagawara A (2004) Polo-like kinase 1 (Plk1) inhibits p53 function by physical interaction and phosphorylation. The Journal of biological chemistry 279: 25549–25561

Barr FA, Sillje HH, Nigg EA (2004) Polo-like kinases and the orchestration of cell division. Nature reviews Molecular cell biology 5: 429–440

Ben-Bassat H, Goldblum N, Mitrani S, Goldblum T, Yoffey JM, Cohen MM, Bentwich Z, Ramot B, Klein E, Klein G (1977) Establishment in continuous culture of a new type of lymphocyte from a “Burkitt like” malignant lymphoma (line D.G.-75). Int J Cancer 19: 27–33

Caduff N, McHugh D, Murer A, Rämer P, Raykova A, Landtwing V, Rieble L, Keller CW, Prummer M, Hoffmann L et al (2020) Immunosuppressive FK506 treatment leads to more frequent EBV-associated lymphoproliferative disease in humanized mice. PLoS pathogens 16: e1008477

Chabot PR, Raiola L, Lussier-Price M, Morse T, Arseneault G, Archambault J, Omichinski JG (2014) Structural and functional characterization of a complex between the acidic transactivation domain of EBNA2 and the Tfb1/p62 subunit of TFIIH. PLoS pathogens 10: e1004042

Cheng K-Y, Lowe ED, Sinclair J, Nigg EA, Johnson LN (2003) The crystal structure of the human polo-like kinase-1 polo box domain and its phospho-peptide complex. The EMBO journal 22: 5757–5768

Cohen JI (1992) A region of herpes simplex virus VP16 can substitute for a transforming domain of Epstein-Barr virus nuclear protein 2. Proceedings of the National Academy of Sciences of the United States of America 89: 8030–8034

Cohen JI, Kieff E (1991) An Epstein-Barr virus nuclear protein 2 domain essential for transformation is a direct transcriptional activator. Journal of virology 65: 5880–5885.

Cohen JI, Wang F, Kieff E (1991) Epstein-Barr virus nuclear protein 2 mutations define essential domains for transformation and transactivation. Journal of virology 65: 2545–2554

Cox J, Mann M (2008) MaxQuant enables high peptide identification rates, individualized p.p.b.-range mass accuracies and proteome-wide protein quantification. Nature biotechnology 26: 1367–1372

de Carcer G (2019) The Mitotic Cancer Target Polo-Like Kinase 1: Oncogene or Tumor Suppressor? Genes 10

Elia AE, Cantley LC, Yaffe MB (2003a) Proteomic screen finds pSer/pThr-binding domain localizing Plk1 to mitotic substrates. Science 299: 1228–1231

Elia AE, Rellos P, Haire LF, Chao JW, Ivins FJ, Hoepker K, Mohammad D, Cantley LC, Smerdon SJ, Yaffe MB (2003b) The molecular basis for phosphodependent substrate targeting and regulation of Plks by the Polo-box domain. Cell 115: 83–95

Farrell CJ, Lee JM, Shin EC, Cebrat M, Cole PA, Hayward SD (2004) Inhibition of Epstein-Barr virus-induced growth proliferation by a nuclear antigen EBNA2-TAT peptide. Proceedings of the National Academy of Sciences of the United States of America 101: 4625–4630

Farrell PJ (2019) Epstein-Barr Virus and Cancer. Annual review of pathology 14: 29–53

Francis MS, Amer AAA, Milton DL, Costa TRD (2017) Site-Directed Mutagenesis and Its Application in Studying the Interactions of T3S Components. In: Type 3 Secretion Systems: Methods and Protocols, Nilles M.L., Condry D.L.J. (eds.) pp. 11–31. Springer New York: New York, NY

Friberg A, Thumann S, Hennig J, Zou P, Nossner E, Ling PD, Sattler M, Kempkes B (2015) The EBNA-2 N-Terminal Transactivation Domain Folds into a Dimeric Structure Required for Target Gene Activation. PLoS pathogens 11: e1004910

Gheghiani L, Loew D, Lombard B, Mansfeld J, Gavet O (2017) PLK1 Activation in Late G2 Sets Up Commitment to Mitosis. Cell Rep 19: 2060–2073

Glaser LV, Rieger S, Thumann S, Beer S, Kuklik-Roos C, Martin DE, Maier KC, Harth-Hertle ML, Gruning B, Backofen R et al (2017) EBF1 binds to EBNA2 and promotes the assembly of EBNA2 chromatin complexes in B cells. PLoS pathogens 13: e1006664

Harter MR, Liu CD, Shen CL, Gonzalez-Hurtado E, Zhang ZM, Xu M, Martinez E, Peng CW, Song J (2016) BS69/ZMYND11 C-Terminal Domains Bind and Inhibit EBNA2. PLoS pathogens 12: e1005414

Henkel T, Ling PD, Hayward SD, Peterson MG (1994) Mediation of Epstein-Barr virus EBNA2 transactivation by recombination signal-binding protein J kappa. Science 265: 92–95

Hsieh JJ, Hayward SD (1995) Masking of the CBF1/RBPJ kappa transcriptional repression domain by Epstein-Barr virus EBNA2. Science 268: 560–563

Kempkes B, Ling PD (2015) EBNA2 and Its Coactivator EBNA-LP. Current topics in microbiology and immunology 391: 35–59

Koida N, Ozaki T, Yamamoto H, Ono S, Koda T, Ando K, Okoshi R, Kamijo T, Omura K, Nakagawara A (2008) Inhibitory role of Plk1 in the regulation of p73-dependent apoptosis through physical interaction and phosphorylation. The Journal of biological chemistry 283: 85558563

Kuppers R, Schneider M, Hansmann ML (2019) Laser-Based Microdissection of Single Cells from Tissue Sections and PCR Analysis of Rearranged Immunoglobulin Genes from Isolated Normal and Malignant Human B Cells. Methods Mol Biol 1956: 61–75

Kuppers R, Zhao M, Rajewsky K, Hansmann ML (1993) Detection of clonal B cell populations in paraffin-embedded tissues by polymerase chain reaction. Am J Pathol 143: 230–239

Kwiatkowski B, Chen SY, Schubach WH (2004) CKII site in Epstein-Barr virus nuclear protein 2 controls binding to hSNF5/Ini1 and is important for growth transformation. Journal of virology 78: 6067–6072

Lee KH, Hwang J-A, Kim S-O, Kim JH, Shin SC, Kim EE, Lee KS, Rhee K, Jeon BH, Bang JK et al (2018) Phosphorylation of human enhancer filamentation 1 (HEF1) stimulates interaction with Polo-like kinase 1 leading to HEF1 localization to focal adhesions. The Journal of biological chemistry 293: 847–862

Lee KS, Park JE, Kang YH, Kim TS, Bang JK (2014) Mechanisms underlying Plk1 polo-box domain-mediated biological processes and their physiological significance. Molecules and cells 37: 286–294

Ling PD, Hayward SD (1995) Contribution of conserved amino acids in mediating the interaction between EBNA2 and CBF1/RBPJk. Journal of virology 69: 1944–1950.

Liu Z, Sun Q, Wang X (2017) PLK1, A Potential Target for Cancer Therapy. Translational oncology 10: 22–32

Longnecker RM, Kieff E, Cohen JI (2013) Epstein-Barr virus. In: Fields Virology, Knipe D.M., Howley P.M., Cohen J.I., Griffin D.E., Lamb R.A., Martin M.A., Racianello V.R., Roizman B. (eds.) pp. 1898–1959. Lippincott Williams and Wilkins: Philadelphia

Lowery DM, Lim D, Yaffe MB (2005) Structure and function of Polo-like kinases. Oncogene 24: 248–259

Lu F, Chen HS, Kossenkov AV, DeWispeleare K, Won KJ, Lieberman PM (2016) EBNA2 Drives Formation of New Chromosome Binding Sites and Target Genes for B-Cell Master Regulatory Transcription Factors RBP-jkappa and EBF1. PLoS pathogens 12: e1005339

Marceau AH, Brison CM, Nerli S, Arsenault HE, McShan AC, Chen E, Lee HW, Benanti JA, Sgourakis NG, Rubin SM (2019) An order-to-disorder structural switch activates the FoxM1 transcription factor. eLife 8

Martin BT, Strebhardt K (2006) Polo-like kinase 1: target and regulator of transcriptional control. Cell cycle 5: 2881–2885

Minoguchi S, Taniguchi Y, Kato H, Okazaki T, Strobl LJ, Zimber-Strobl U, Bornkamm GW, Honjo T (1997) RBP-L, a transcription factor related to RBP-Jkappa. Molecular and cellular biology 17: 2679–2687

Mrozek-Gorska P, Buschle A, Pich D, Schwarzmayr T, Fechtner R, Scialdone A, Hammerschmidt W (2019) Epstein-Barr virus reprograms human B lymphocytes immediately in the prelatent phase of infection. Proceedings of the National Academy of Sciences of the United States of America 116: 16046–16055

Munz C (2019) Latency and lytic replication in Epstein-Barr virus-associated oncogenesis. Nat Rev Microbiol 17: 691–700

Nakajima H, Toyoshima-Morimoto F, Taniguchi E, Nishida E (2003) Identification of a consensus motif for Plk (Polo-like kinase) phosphorylation reveals Myt1 as a Plk1 substrate. The Journal of biological chemistry 278: 25277–25280

Pavlovsky AG, Liu X, Faehnle CR, Potente N, Viola RE (2012) Structural characterization of inhibitors with selectivity against members of a homologous enzyme family. Chem Biol Drug Des 79: 128–136

Pich D, Mrozek-Gorska P, Bouvet M, Sugimoto A, Akidil E, Grundhoff A, Hamperl S, Ling PD, Hammerschmidt W (2019) First Days in the Life of Naive Human B Lymphocytes Infected with Epstein-Barr Virus. MBio 10

Ponnusamy R, Khatri R, Correia PB, Wood CD, Mancini EJ, Farrell PJ, West MJ (2019) Increased association between Epstein-Barr virus EBNA2 from type 2 strains and the transcriptional repressor BS69 restricts EBNA2 activity. PLoS pathogens 15: e1007458

Pulvertaft JV (1964) Cytology of Burkitt’s Tumour (African Lymphoma). Lancet 39: 238–240

Raab M, Sanhaji M, Matthess Y, Horlin A, Lorenz I, Dotsch C, Habbe N, Waidmann O, Kurunci-Csacsko E, Firestein R et al (2018) PLK1 has tumor-suppressive potential in APC-truncated colon cancer cells. Nature communications 9: 1106

Rajewsky K (1996) Clonal selection and learning in the antibody system. Nature 381: 751–758

Ren Y, Bi C, Zhao X, Lwin T, Wang C, Yuan J, Silva AS, Shah BD, Fang B, Li T et al (2018) PLK1 stabilizes a MYC-dependent kinase network in aggressive B cell lymphomas. The Journal of clinical investigation 128: 5517–5530

Rodel F, Zhou S, Gyorffy B, Raab M, Sanhaji M, Mandal R, Martin D, Becker S, Strebhardt K (2020) The Prognostic Relevance of the Proliferation Markers Ki-67 and Plk1 in Early-Stage Ovarian Cancer Patients With Serous, Low-Grade Carcinoma Based on mRNA and Protein Expression. Front Oncol 10: 558932

Rosenblum D, Gutkin A, Kedmi R, Ramishetti S, Veiga N, Jacobi AM, Schubert MS, Friedmann-Morvinski D, Cohen ZR, Behlke MA et al (2020) CRISPR-Cas9 genome editing using targeted lipid nanoparticles for cancer therapy. Science Advances 6: eabc9450

Shannon-Lowe C, Rickinson A (2019) The Global Landscape of EBV-Associated Tumors. Front Oncol 9: 713

Shevchenko A, Wilm M, Vorm O, Mann M (1996) Mass spectrometric sequencing of proteins silver-stained polyacrylamide gels. Anal Chem 68: 850–858

Shultz LD, Saito Y, Najima Y, Tanaka S, Ochi T, Tomizawa M, Doi T, Sone A, Suzuki N, Fujiwara H et al (2010) Generation of functional human T-cell subsets with HLA-restricted immune responses in HLA class I expressing NOD/SCID/IL2r gamma(null) humanized mice. Proceedings of the National Academy of Sciences of the United States of America 107: 13022–13027

Śledź P, Lang S, Stubbs CJ, Abell C (2012) High-throughput interrogation of ligand binding mode using a fluorescence-based assay. Angewandte Chemie (International ed in English) 51: 7680–7683

Strebhardt K (2010) Multifaceted polo-like kinases: drug targets and antitargets for cancer therapy. Nat Rev Drug Discov 9: 643–660

Strebhardt K, Ullrich A (2006) Targeting polo-like kinase 1 for cancer therapy. Nat Rev Cancer 6: 321–330

Strowig T, Gurer C, Ploss A, Liu Y-F, Arrey F, Sashihara J, Koo G, Rice CM, Young JW, Chadburn A et al (2009a) Priming of protective T cell responses against virus-induced tumors in mice with human immune system components. The Journal of experimental medicine 206: 1423–1434

Strowig T, Gurer C, Ploss A, Liu YF, Arrey F, Sashihara J, Koo G, Rice CM, Young JW, Chadburn A et al (2009b) Priming of protective T cell responses against virus-induced tumors in mice with human immune system components. The Journal of experimental medicine 206: 1423–1434

Thorley-Lawson DA, Gross A (2004) Persistence of the Epstein-Barr virus and the origins of associated lymphomas. N Engl J Med 350: 1328–1337

Tong X, Drapkin R, Reinberg D, Kieff E (1995a) The 62- and 80-kDa subunits of transcription factor IIH mediate the interaction with Epstein-Barr virus nuclear protein 2. Proceedings of the National Academy of Sciences of the United States of America 92: 3259–3263.

Tong X, Drapkin R, Yalamanchili R, Mosialos G, Kieff E (1995b) The Epstein-Barr virus nuclear protein 2 acidic domain forms a complex with a novel cellular coactivator that can interact with TFIIE. Molecular and cellular biology 15: 4735–4744.

Tong X, Wang F, Thut CJ, Kieff E (1995c) The Epstein-Barr virus nuclear protein 2 acidic domain can interact with TFIIB, TAF40, and RPA70 but not with TATA-binding protein. Journal of virology 69: 585–588.

Vizcaino JA, Cote RG, Csordas A, Dianes JA, Fabregat A, Foster JM, Griss J, Alpi E, Birim M, Contell J et al (2013) The PRoteomics IDEntifications (PRIDE) database and associated tools: status in 2013. Nucleic acids research 41: D1063–1069

Vizcaino JA, Csordas A, Del-Toro N, Dianes JA, Griss J, Lavidas I, Mayer G, Perez-Riverol Y, Reisinger F, Ternent T et al (2016) 2016 update of the PRIDE database and its related tools. Nucleic acids research 44: 11033

Wang L, Grossman SR, Kieff E (2000) Epstein-Barr virus nuclear protein 2 interacts with p300, CBP, and PCAF histone acetyltransferases in activation of the LMP1 promoter. Proceedings of the National Academy of Sciences of the United States of America 97: 430–435.

Wang S, Zhao Y, Leiby M, Zhu J (2009) A new positive/negative selection scheme for precise BAC recombineering. Mol Biotechnol 42: 110–116

Warming S, Costantino N, Court DL, Jenkins NA, Copeland NG (2005) Simple and highly efficient BAC recombineering using galK selection. Nucleic acids research 33: e36

Watanabe N, Arai H, Iwasaki J, Shiina M, Ogata K, Hunter T, Osada H (2005) Cyclin-dependent kinase (CDK) phosphorylation destabilizes somatic Wee1 via multiple pathways. Proceedings of the National Academy of Sciences of the United States of America 102: 11663–11668

West MJ (2017) Chromatin reorganisation in Epstein-Barr virus-infected cells and its role in cancer development. Current opinion in virology 26: 149–155

Wolf G, Elez R, Doermer A, Holtrich U, Ackermann H, Stutte HJ, Altmannsberger H-M, Rübsamen-Waigmann H, Strebhardt K (1997) Prognostic significance of polo-like kinase (PLK) expression in non-small cell lung cancer. Oncogene 14: 543–549

Yuan C, Wang L, Zhou L, Fu Z (2014) The function of FOXO1 in the late phases of the cell cycle is suppressed by PLK1-mediated phosphorylation. Cell cycle 13: 807–819

Yuan J, Eckerdt F, Bereiter-Hahn J, Kurunci-Csacsko E, Kaufmann M, Strebhardt K (2002) Cooperative phosphorylation including the activity of polo-like kinase 1 regulates the subcellular localization of cyclin B1. Oncogene 21: 8282–8292

Yuan J, Horlin A, Hock B, Stutte HJ, Rubsamen-Waigmann H, Strebhardt K (1997) Polo-like kinase, a novel marker for cellular proliferation. Am J Pathol 150: 1165–1172

Yue W, Davenport MG, Shackelford J, Pagano JS (2004) Mitosis-specific hyperphosphorylation of Epstein-Barr virus nuclear antigen 2 suppresses its function. Journal of virology 78: 3542–3552

Yue W, Gershburg E, Pagano JS (2005) Hyperphosphorylation of EBNA2 by Epstein-Barr virus protein kinase suppresses transactivation of the LMP1 promoter. Journal of virology 79: 5880–5885

Yue W, Shackelford J, Pagano JS (2006) cdc2/cyclin B1-dependent phosphorylation of EBNA2 at Ser243 regulates its function in mitosis. Journal of virology 80: 2045–2050

Yun S-M, Moulaei T, Lim D, Bang JK, Park J-E, Shenoy SR, Liu F, Kang YH, Liao C, Soung N-K et al (2009) Structural and functional analyses of minimal phosphopeptides targeting the polobox domain of polo-like kinase 1. Nature structural & molecular biology 16: 876–882

Zdimerova H, Murer A, Engelmann C, Raykova A, Deng Y, Gujer C, Rühl J, McHugh D, Caduff N, Naghavian R et al (2020) Attenuated immune control of Epstein-Barr virus in humanized mice is associated with the multiple sclerosis risk factor HLA-DR15. European journal of immunology

